# miRNA-mediated inhibition of an actomyosin network in hippocampal pyramidal neurons restricts sociability in adult male mice

**DOI:** 10.1101/2023.11.08.566242

**Authors:** Ramanathan Narayanan, Brunno Rocha Levone, Jochen Winterer, Prakruti Nanda, Alexander Mueller Ranft, Roberto Fiore, Pierre-Luc Germain, Marija Mihailovich, Giuseppe Testa, Gerhard Schratt

## Abstract

Social deficits are frequently observed in patients suffering from neurodevelopmental disorders, but the molecular mechanisms regulating sociability are still poorly understood. We recently reported that the loss of the microRNA cluster miR-379-410 leads to hypersocial behavior and anxiety in mice. Here, we show that ablating miR-379-410 in excitatory neurons of the postnatal mouse hippocampus recapitulates hypersociability, but not anxiety. At the cellular level, miR-379-410 loss in excitatory neurons leads to increased excitatory synaptic transmission and upregulation of an actomyosin gene network. Re-expression of three cluster miRNAs, as well as pharmacological inhibition of the actomyosin activator ROCK, was sufficient to reinstate normal sociability in miR-379-410 knockout mice. Several actomyosin genes and miR-379-410 family members were reciprocally dysregulated in isogenic human iPSC-derived neurons harboring a deletion present in Williams-Beuren-Syndrome patients, which are characterized by hypersocial behavior. Together, our results unveil a novel microRNA-actomyosin pathway involved in the control of sociability.

## Introduction

Social interactions are fundamental and adaptive components of the biology of numerous species, including rodents and humans. Over the course of mouse development, sociability is successively reduced in particular in males at the expense of more aggressive behavior which underlies the development of dominance-submission relationships within the social group [1–3]. Sociability has a high variability in the human population, with a portion of people outside the normal range [4]. While psychiatric disorders (e.g., schizophrenia) and autism spectrum disorders (ASD) are typically associated with a deficit in social behavior, the opposite trait of hypersociability is exhibited by individuals with specific neurodevelopmental disorders, e.g., Angelman Syndrome (AS) and Williams-Beuren Syndrome (WBS). Together, these observations suggest the existence of a yet unknown neural mechanisms that restrict sociability in a developmental stage dependent manner.

Neural circuits for social interactions have been described in great detail using opto- and chemogenetic approaches and include connections between the medial prefrontal cortex (mPFC), hippocampus (HPC), basolateral amygdala (BLA), nucleus accumbens (NAc) and anterior cingulate cortex (ACC), and other brain regions [5, 6]. The hippocampus in particular plays important roles in the modulation of social interaction, preference, and recognition memory [7]. The BLA inputs to CA1 pyramidal neurons within the ventral HPC bi-directionally modulating social interaction and anxiety-related behavior [8, 9]. In contrast, projections from ventral CA1 neurons to the NAc shell play a necessary and sufficient role in social memory [10]. In addition, dorsal hippocampal CA2 and CA3 regions have been implicated in social memory [11, 12].

Important insights regarding the molecular pathways involved in the control of social interaction came from genetic studies identifying ASD risk genes, which highlight a positive role for glutamatergic synaptic transmission, gene expression regulation and neuromodulatory peptides (such as Oxytocin and Vasopressin) in sociability [SPARK gene list; SPARKforAutism.org]. For example, deficiency in several components of glutamatergic synapses, including adhesion molecules (Neuroligins and Neurexins), scaffolding proteins (Shank, Syngap), receptors (Grin1, 2) and structural proteins (actomyosin cytoskeleton) impair social functioning [13]. In addition, perturbance of gene regulatory pathways at the level of transcription (e.g., chromatin complexes) [14] and translation (e.g., mTOR pathway) [15] has been repeatedly implicated in social dysfunction. In contrast, relatively little is known regarding mechanisms that naturally restrict sociability and whose inactivation leads to hypersociability.

Knowledge about such pathways could help to devise strategies to promote sociability in neuropsychiatric disorders. In WBS, social deficits have been mapped to the deletion of the related *Gtf2i* and *Gtf2ird1* genes, which encode transcription factors, using human genetic and mouse behavioral approaches [16]. AS is caused by loss-of-function mutations in the *Ube3a* gene, which encodes a ubiquitin ligase with roles in transcriptional control and synaptic protein homeostasis. In addition, a handful of other loss-of-function mutations leads to hypersociability in mouse models, whereby the underlying molecular mechanisms are poorly understood [4].

Due to their well-documented role in the development and plasticity of neural circuits, such as the hippocampus, microRNAs (miRNAs) represent strong candidates for regulators of social interactions. miRNAs are a large class of non-coding RNAs which act as post-transcriptional repressors of gene expression [17]. The maternally imprinted miR379-410 cluster represents the largest mammalian-specific miRNA cluster harboring 38 miRNA genes [18]. Maternal deletion of miR379-410 in induced mouse neurons results in elevated excitatory synaptic transmission [19]. miR-134-5p, the most extensively studied miR379-410 member, inhibits dendritic spine morphogenesis and synaptic function in rat hippocampal neurons by repressing the synthesis of Limk1, a kinase controlling actin dynamics and deleted in WBS [20]. Furthermore, dendritic activity of several miR379-410 miRNAs is controlled by a non-coding RNA originating from the AS gene Ube3a [21]. Together, these previous studies implicate miR379-410 in the regulation of synaptic transmission in the context of hypersociability disorders.

We recently reported that ubiquitous loss of miR379-410 leads to hypersociability and increased anxiety-related behavior in mice [22]. Behavioral phenotypes were associated with upregulation of glutamatergic synaptic transmission in the hippocampus. However, from this study, several questions remained unresolved: in which tissue and cell type exerts miR379-410 its effect on social behavior and anxiety? What are the functionally important miR379-410 members? What are the exact mechanisms mediating the behavioral alterations? And finally, what are the implications for disorders characterized by hypersociability and anxiety, i.e. WBS?

In the present study, we obtained evidence that three specific miR379-410 cluster members restrict sociability in adult male mice by inhibiting the expression of an actomyosin network in CA1 hippocampal pyramidal neurons. Furthermore, we found that this pathway is disinhibited in human neurons carrying a WBS genetic deletion. We conclude that targeting this novel pathway could have therapeutic potential for WBS and other neurodevelopmental conditions characterized by social impairments.

## Results

### Conditional deletion of miR379-410 in excitatory neurons of the mouse forebrain leads to hypersocial behavior only in adult male mice, but not in females or juvenile mice

Previously, we have shown that CMV-Cre driven ubiquitous deletion of the miR379-410 cluster in mice leads to hypersociability and increased anxiety-like behavior in mice [22]. To address the specific involvement of principal excitatory forebrain neurons, the cell type wherein miR379-410 is most highly expressed [23], in this behavior, we crossed miR379-410 flox mice to a forebrain-specific Cre deleter strain (Emx1-Cre), resulting in conditional miR379-410 knockout (cKO) mice from embryonic day 12.5 throughout adulthood (Suppl. Fig. 1A-B). qPCR analysis of RNA obtained from WT and cKO mice revealed a highly penetrant depletion of the long noncoding RNA Mirg, the miR379-410 host gene (Suppl. Fig. 1C), as well as a reduction in the expression of several mature miR379-410-derived miRNAs, in the hippocampus (Hc) and in the cortex (Cx) of cKO mice (Suppl. Fig. 1D). In contrast, the expression of members of the miR379-410 cluster was not affected in the striatum (St), a brain region where Emx1-Cre is not active (Suppl. Fig. 1E).

To establish whether the genetic deletion of the miR379-410 cluster in excitatory neurons drives changes in behavior, adult cKO and WT mice (PW15) were submitted to a battery of behavioral testing (Fig. 1A). Adult cKO male mice displayed a hypersocial behavior both in the reciprocal social interaction test (Fig. 1B) and in social preference in the three chambers test (Fig. 1C). Hypersociability was exclusively observed in male, but not female cKO mice (Fig. 1B-C). In contrast, no genotype- or sex-dependent effects were observed in social memory (three chambers test (Fig. 1D)), anxiety-like behavior (open field test (Suppl. Fig. 2A), light-dark box (Suppl. Fig. 2B), elevated plus maze (Suppl. Fig. 2C)) or in memory performance (novel object recognition test (Suppl. Fig. 2D), fear conditioning (Suppl. Fig. 2E)). Moreover, sociability, anxiety and memory were indistinguishable between juvenile (PW5) cKO and WT mice (Suppl Fig. 3A-E). This finding is consistent with the low expression of the miR379-410 cluster miRNAs miR-134-5p and miR-485-5p at the early postnatal stage (PW4; Suppl. Fig. 3F). Taken together, our results suggest that miR379-410 function in excitatory forebrain neurons is required for the control of sociability, but is dispensable for anxiety-related behavior and memory in adult male mice.

**Figure 1:**
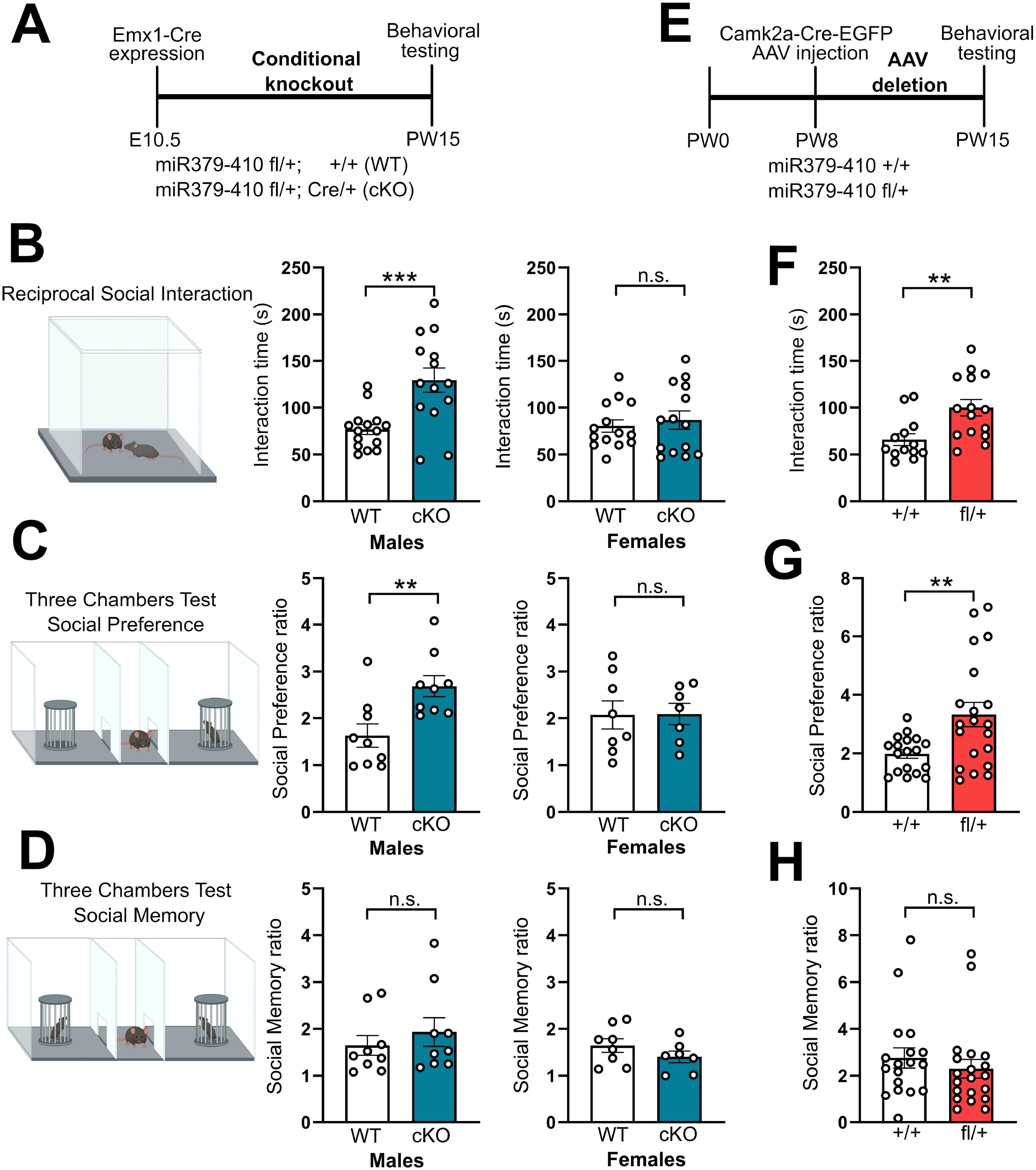
Conditional knockout and acute AAV-mediated deletion of the miR379-410 cluster in excitatory neurons of the forebrain leads to hypersocial behavior in adult mice. **(A)** Timeline of the Emx-1-Cre expression and behavioral testing in the Genetic deletion mouse model. The genetic deletion of the miR379-410 cluster (cKO) induces hypersocial behavior in the Reciprocal Social Interaction test **(B)** (males p = 0.0007, females p = 0.594) and in the Social Preference in the Three Chambers test **(C)** (males p = 0.0064, females p = 0.961), specifically in male (left graphs), but not in female (right graphs) adult mice. **(D)** No changes in Social Memory were observed in the Three Chambers test (males p = 0.455, females p = 0.237), neither in male (left graph), nor in female (right graph) adult mice. **(E)** Timeline of the acute AAV deletion of the miR379-410 cluster in excitatory neurons of the hippocampus and behavioral testing. The acute deletion of the miR379-410 cluster in the hippocampus of adult male mice is enough to induce hypersocial behavior in the Reciprocal Social Interaction test **(F)** (p = 0.0046) and in the Social Preference in the Three Chambers test **(G)** (p = 0.0059), with no changes in Social Memory in the Three Chambers test **(H)** (p = 0.437). Statistics: unpaired t-test.

### Acute AAV-mediated knockout of miR379-410 in hippocampal pyramidal neurons of adult mice leads to hypersocial behavior

We sought to further investigate the specific brain regions and neuronal cell types involved in miR379-410 regulated social behavior. We focused on the hippocampus due to the reported functions of miR379-410 members in hippocampal pyramidal neurons *in vitro* [24, 25] and the well-established role of the hippocampus in social behavior. We injected into the hippocampus of adult miR379-410 flox or WT control mice (Figure 1E and Suppl Fig. 4A) a recombinant adeno-associated virus (rAAV) expressing Cre under the control of the excitatory-neuron specific Camk2a promoter (AAV-Camk2a-Cre-eGFP). This led to a highly penetrant and specific Cre-eGFP expression in CA1-3 principal layers of the hippocampus (Suppl. Fig. 4B). While the expression level of an unrelated miRNA (miR-132-3p) was unchanged, hippocampal levels of miR379-410 members (miR-134-5p and miR-485-5p) were reduced by about 50% (Suppl. Fig. 4C), demonstrating successful recombination of the floxed miR379-410 allele. The reduction in miR379-410 expression specifically in the hippocampus of adult male mice was sufficient to elicit hypersocial behavior compared to WT controls, as evidenced both by an increase in reciprocal social interaction (Fig. 1F) and in social preference in the three chambers test (Fig. 1G). Similar to the Emx1-Cre driven miR379-410 knockout, the Camk2a-Cre driven ablation of the cluster had no effects in social memory (Fig. 1H), in anxiety-related behavior and nor in memory performance (Suppl. Fig. 4D-F). Thus, miR379-410 expression in hippocampal pyramidal neurons is required to restrict sociability in adult male mice.

### miR379-410 deficiency in hippocampal pyramidal neurons of adult male mice leads to increased excitatory synaptic transmission

To investigate potential synaptic alterations underlying the hypersociability phenotype of miR379-410 knockout mice, we performed patch-clamp electrophysiological recordings in hippocampal slices from PW15 WT versus cKO male mice. Consistent with previous results, whole cell patch-clamp recordings revealed a significant enhancement of mEPSC amplitudes, but not frequency, in CA1 pyramidal neurons from cKO compared to WT mice (Fig. 2A-C). Paired-pulse facilitation (PPF) was not affected in cKO slices, arguing against a presynaptic deficit (Suppl. Fig. 5A). Furthermore, we similarly observed an increase in mEPSC amplitude, but not frequency, in CA1 pyramidal neurons from miR379-410 flox mice injected with AAV-Camk2a-Cre-eGFP (Fig. 2D-F). mEPSC amplitudes, but not frequency, further positively correlated with the social preference index in mice injected with AAV-Camk2a-Cre-eGFP (Suppl. Fig. 5B-C). Thus, miR379-410 mediated inhibition of sociability coincides with an inhibition of excitatory synaptic transmission onto hippocampal pyramidal neurons.

**Figure 2:**
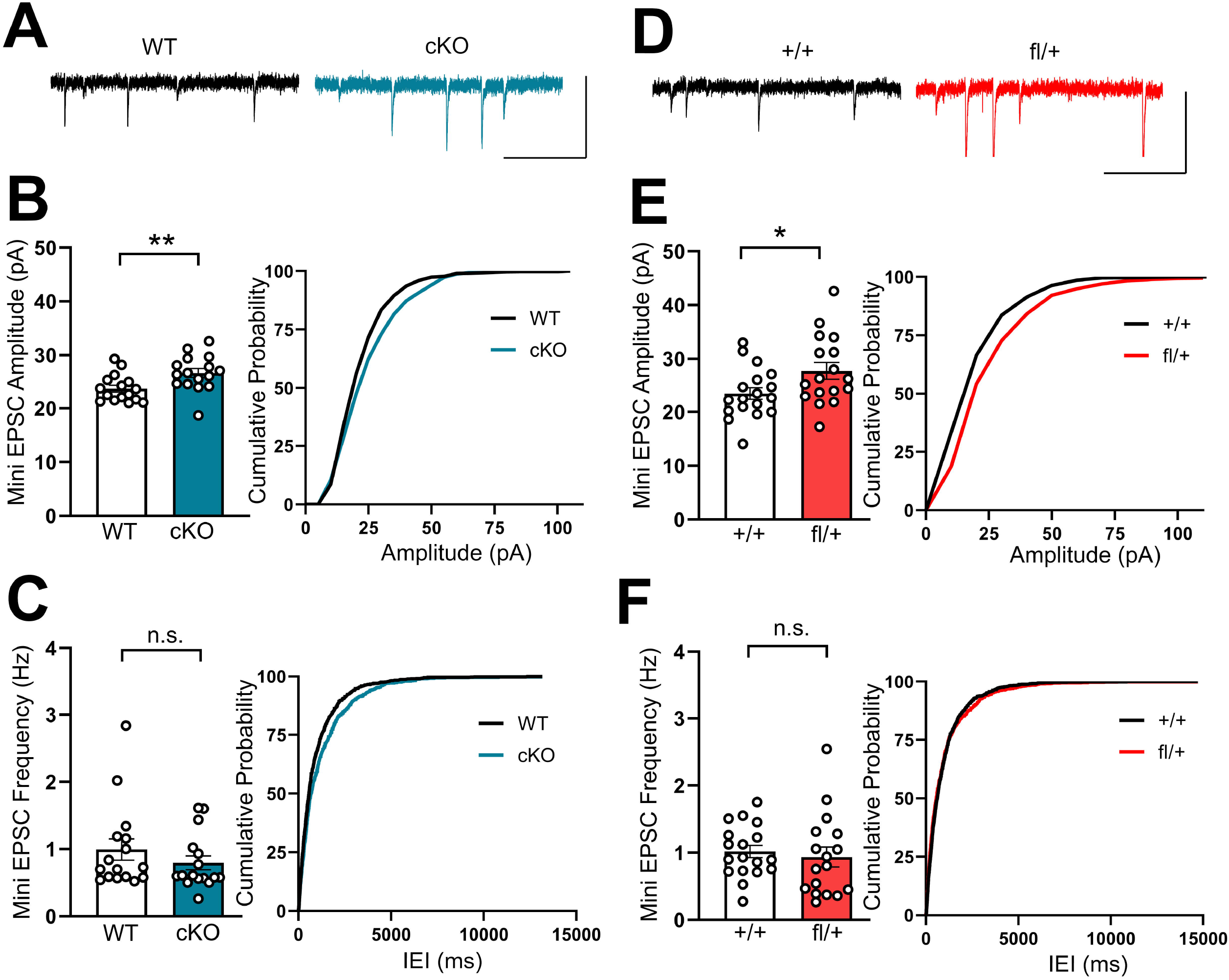
Conditional knockout and acute AAV-mediated deletion of the miR379-410 cluster increases excitatory synaptic transmission in hippocampal pyramidal neurons. **(A)** Representative mini EPSC traces in WT and miR379-410 knockout mouse hippocampus brain slices. The genetic deletion of the miR379-410 cluster in excitatory neurons increases the mini EPSC amplitude **(B)** (p = 0.0083) in hippocampal brain slices, with no change in mini EPSC frequency **(C)** (p = 0.303). This increase in mini EPSC frequency was also observed in the acute AAV deletion of the cluster **(D, E)** (p = 0.032), with no changes in mini EPSC frequency **(F)** (p = 0.635). Statistics: unpaired t-test.

### Loss of miR379-410 in adult mouse hippocampus leads to the induction of an actomyosin gene network

To obtain insight into the molecular mechanisms underlying hypersociability and enhanced excitatory synaptic transmission upon miR379-410 ablation, we performed polyA-seq of total RNA obtained from the hippocampus of cKO (Emx1-Cre) and WT control mice. Differential gene expression (DEG) analysis identified a total of 76 differentially expressed genes (37 up, 39 down; FDR<0.05; Fig. 3A-B). Gene set enrichment analysis revealed an expected upregulation of the actomyosin gene complex (Fig. 3C), including a network of several actomyosin genes (Myh11, Myl9, Acta2, Tagln, Cnn1, Tpm2, Mustn1, Suppl. Fig. 6A).

**Figure 3:**
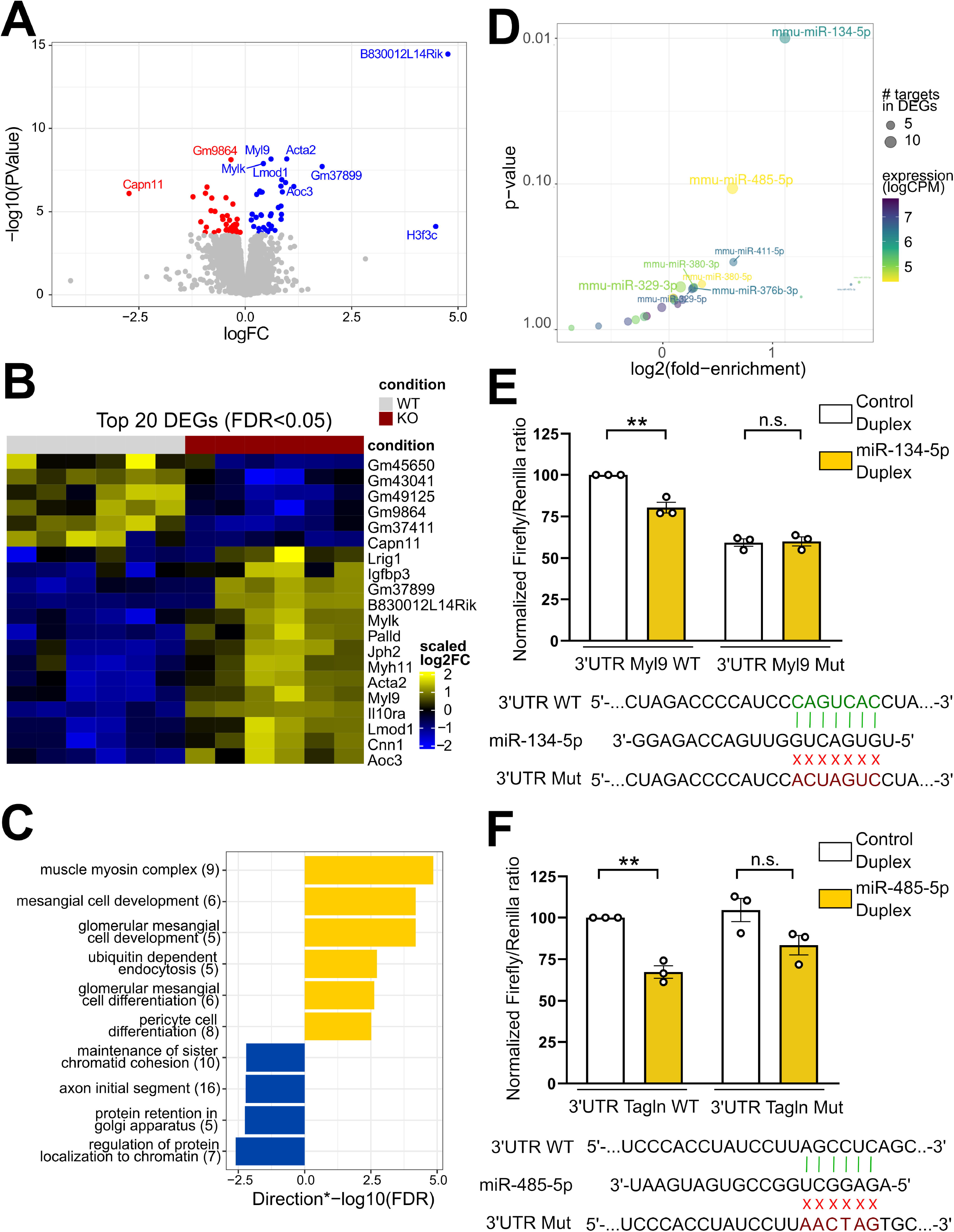
Conditional knockout of the miR379-410 cluster leads to the upregulation of an actomyosin gene network in the adult hippocampus. **(A)** Volcano plot of the differential expression in cKO mice (genes with FDR<0.05 are colored, and the top 10 genes are labeled). **(B)** Heatmap of the top 20 differentially expressed genes in cKO mice. **(C)** GSEA analysis reveals that top upregulated genes belong to a muscle myosin complex. **(D)** enrichMiR results on the upregulated genes, restricting to miR379-410 cluster miRNAs. The labeled miRNAs were selected on the basis of high expression, number of targets among DEGs, and enrichment. **(E, F)** Luciferase assays revealed that the expression of a reporter construct containing Myl 3’UTR (E) is reduced upon transfection with miR-134 duplex (resulting in an overexpression of the mature miRNA), (p = 0.0037) and the expression of a reporter construct containing Tagln 3’UTR (F) is reduced upon transfection with miR-485 (p = 0.001). A reporter construct containing Myl9 3’UTR or Tagln 3’UTR with a mutated miR-134 or miR-485 binding site, respectively, did not respond to the duplex transfection (Myl9 p = 0.848, Tagln p = 0.081). Statistics: unpaired t-test (E-F).

Next, we sought to identify the specific miR379-410 miRNAs involved in the control of sociability. We reasoned that binding sites for active miRNAs might be overrepresented in transcripts which are upregulated in the hippocampus of cKO mice. Therefore, we employed our recently developed EnrichMiR bioinformatic tool [26] to compute target enrichment scores for the top most expressed 30% miR379-410 members in the mouse brain (Fig. 3D). Using a cut-off of p<0.05, this analysis identified miR-134-5p as the only miRNA whose binding sites are significantly enriched in upregulated transcript. miR-134-5p has previously been shown to control dendritic spine development, synaptic plasticity and memory [20, 24, 27]. Among the top 10 enriched miRNAs were furthermore miR-329-5p and miR-485-5p, members of miRNA families which have been previously implicated in synapse development and plasticity [25, 28]. Based on this analysis, we decided to include miR-134-5p, miR-485-5p and miR-329-5p for our further functional studies.

When inspecting 3’UTR regions of upregulated actomyosin genes, we observed a high density of high-affinity miR-134-5p, -485-5p and -329-5p binding sites within the corresponding transcripts (Suppl. Fig. 6B). Direct functional interactions were subsequently validated for miR-134-5p/Myl9 and miR-485/Tagln using luciferase reporter assays in rat primary cortical neurons (Fig. 3E-F). Whereas Myl9 and Tagln reporter containing cognate miR binding sites were significantly inhibited by miR-134 and miR-485 mimic transfection, respectively, corresponding miR binding site mutants were not altered in their expression (Fig. 3E-F). Myl9 encodes for a regulatory myosin light chain which plays an important role in the regulation of actomyosin contractility in muscle cells [29]. Using qPCR, we found that Myl9 was expressed in mixed primary rat hippocampal neuron cultures (Suppl. Fig. 6C). The majority of detected Myl9 transcripts originates from neurons, since treatment of the cultures with FUDR, which blocks the proliferation of non-neuronal cell types (e.g., astrocytes), did not reduce Myl9 abundance. Thus, we conclude that at least some of the identified upregulated actomyosin genes are direct targets of miR379-410 in neurons.

### Expression of miR-134/485/329-5p in excitatory neurons of the hippocampus rescues hypersocial behavior of miR379-410 deficient mice and is sufficient to induce hyposocial behavior

To assess whether loss of miR-134-5p, -485-5p and -329-5p is causally involved in the development of hypersocial behavior, we decided to restore their expression in the hippocampus of miR379-410-deficient mice (cKO) (Fig. 4A). Therefore, an rAAV expressing corresponding miR-hairpin (miR-Hp) or control (Ctl-Hp) precursors under the control of the Camk2a promoter were injected into the hippocampus of WT and cKO male mice at PW8, followed by behavioral testing at PW15 (Fig. 4A). Using this approach, miR-485-5p levels were restored to nearly WT levels, whereas a 7-fold and 6-fold upregulation of miR-134-5p and miR-329-5p were observed, respectively (Suppl. Fig. 7A, Suppl. Fig. 8A). No change was observed in the expression of a non-cluster miRNA, used as negative control (miR-129-5p, Suppl. Fig. 7A). Importantly, expression of miR-HP in the adult hippocampus of cKO mice completely rescued sociability in the reciprocal interaction and social preference in the three chambers test (Fig. 4B-C), without affecting social memory (Fig. 4D) or anxiety-like behavior in the open field test (Suppl. Fig. 7B). We then went on to test whether expression of the miR-HP also affects social behavior in WT mice. Compared to the injection of Ctl-HP, miR-HP led to a significant reduction in reciprocal social interaction and social preference in the three chambers test (Fig. 4E-F), without affecting social memory in the three chambers test (Fig. 4G), anxiety-like behavior in the open field test (Suppl. Fig. 8B), light-dark box (Suppl. Fig. 8C) and elevated plus maze (Suppl. Fig. 8D) nor memory performance in the novel object recognition (Suppl. Fig. 8E) and fear conditioning test (Suppl. Fig. 8F). The efficiency and specificity of the miR-HP construct were also further validated in the context of dendritic spine morphogenesis in rat hippocampal neurons *in vitro* (Suppl. Fig. 9A-D). We thus conclude that the expression of a three-miRNA signature (miR-134-5p, miR-485-5p, miR-329-5p) in principal neurons of the hippocampus is necessary and sufficient to restrict sociability in adult male mice.

**Figure 4:**
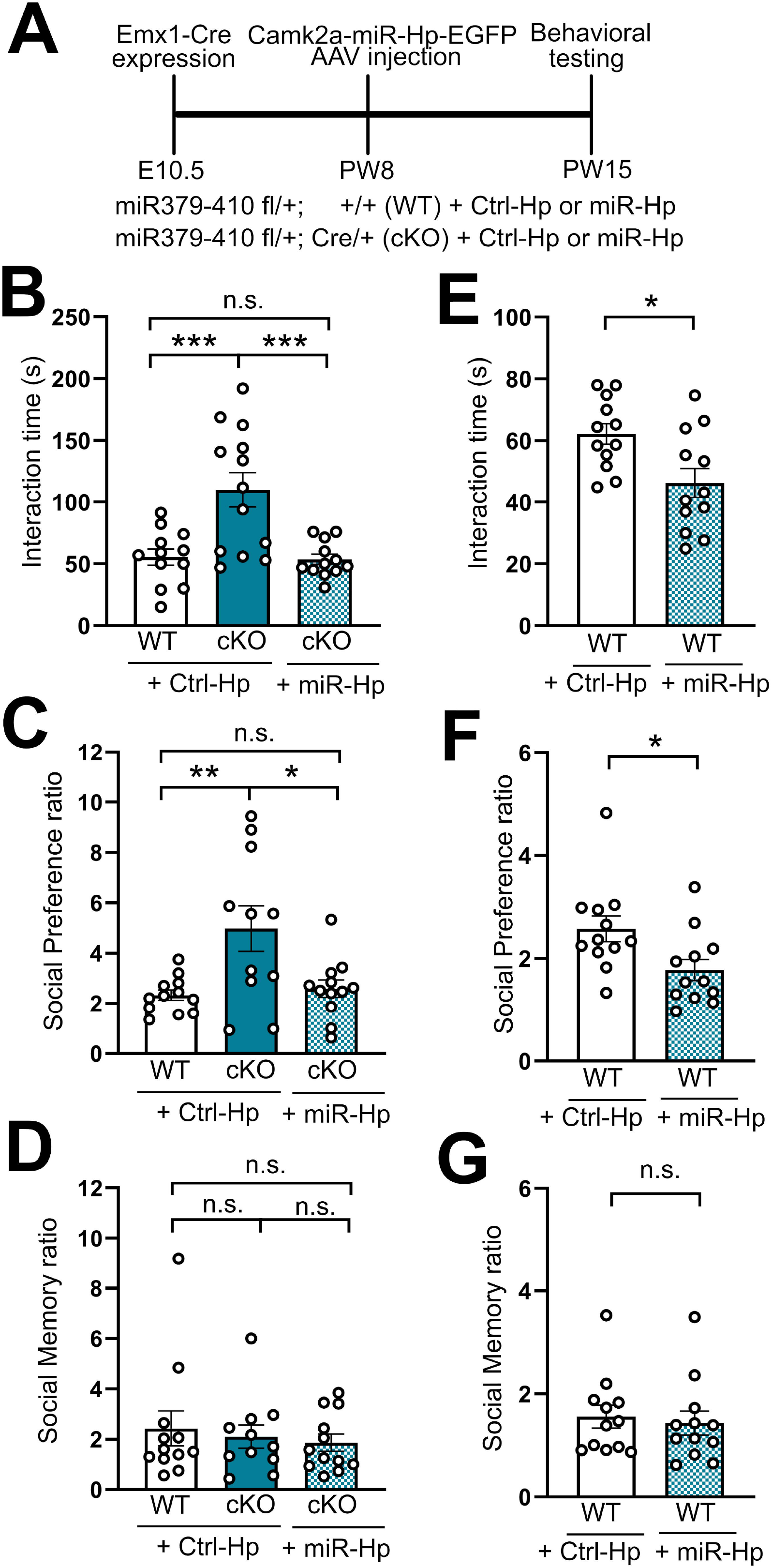
Overexpression of three miR379-410 cluster miRNAs – miR-134-5p, miR-329-5p and miR-485-5p – rescues hypersocial behavior in miR379-410 cKO mice. **(A)** Timeline of the AAV injection of hairpins expressing the miRNAs and behavioral testing. The re-expression of the three-miRNA signature in excitatory neurons of the hippocampus in cKO mice is enough to rescue their hypersocial behavior in the Reciprocal Interaction test **(B)** (WT Ctrl-Hp vs cKO Ctrl-Hp p = 0.011, cKO Ctrl-Hp vs cKO miR-Hp p = 0.065, WT Ctrl-Hp vs cKO miR-Hp p = 0.702) and the Social Preference in the Three Chambers test **(C)** (WT Ctrl-Hp vs cKO Ctrl-Hp p = 0.005, cKO Ctrl-Hp vs cKO miR-Hp p = 0.012, WT Ctrl-Hp vs cKO miR-Hp p = 0.94), with no change in Social Memory in the Three Chambers test **(D)**. The overexpression of these three miRNAs in WT mice reduce sociability in the Reciprocal Interaction test **(E)** (p = 0.011) and the Social Preference in the Three Chambers test **(F)** (p = 0.022), with no change in Social Memory in the Three Chambers test **(G)** (p = 0.707). Statistics: one-way ANOVA with Tukey post-hoc (B-D) and unpaired t-test (E-G).

### Induction of an actomyosin gene network is necessary and sufficient for the regulation of social behavior downstream of miR379-410

As a next step, we explored whether the upregulation of actomyosin genes in miR379-410 knockout mice could play a causal role in hypersociability. Therefore, we first overexpressed two of the genes upregulated in cKO mice, the transcription factor Mustn1 and Myl9, in excitatory neurons of the hippocampus of male mice (PW8) using stereotactic delivery of rAAV-Camk2a-ORF, followed by behavioral testing at PW15 (Fig. 5A, Suppl. Fig. 10A). Mice overexpressing Myl9/Mustn1 (OE) displayed significantly enhanced sociability compared to eGFP-only expressing mice (WT) both in the reciprocal social interaction and in the social preference in the three-chamber test (Fig. 5B-C). Overexpression of Myl9/Mustn1 did not alter social memory (Fig. 5D), anxiety-related behavior in the open field and light-dark box (Suppl. Fig. 10B-C) nor memory performance in the novel object recognition (Suppl. Fig. 10D). Moreover, Myl9/Mustn1 overexpression-driven hypersociability was associated with enhanced excitatory synaptic transmission in CA1 pyramidal neurons as assessed by whole cell patch-clamp recordings in acute hippocampal slices (PW15), in which an increase in mini EPSC amplitude, but not frequency, was observed (Fig. 5E-G).

**Figure 5:**
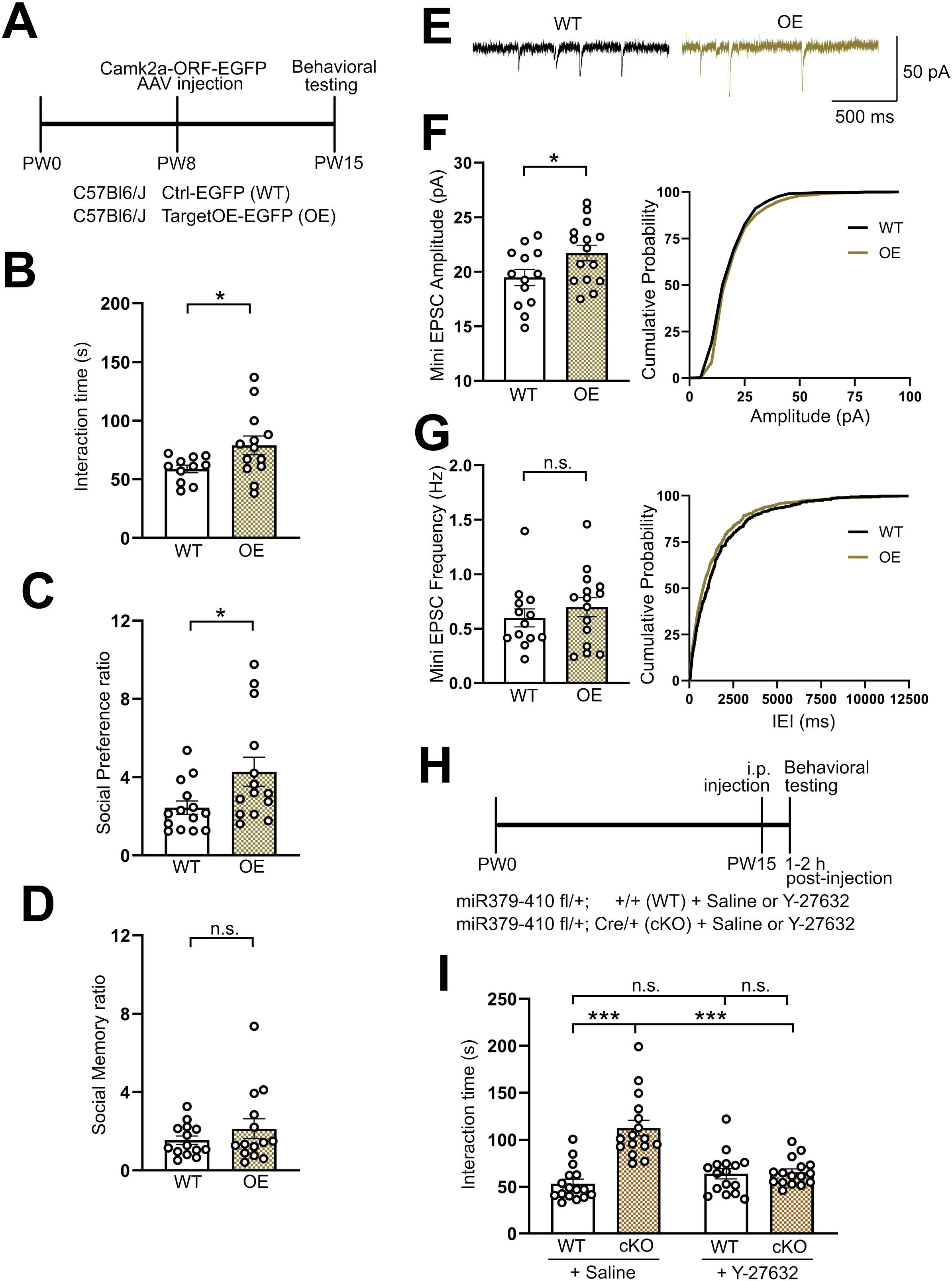
Activation of an actomyosin gene network is necessary and sufficient for hypersocial behavior. **(A)** Timeline of AAV injection for the overexpression of Myl9 and Mustn1 and behavioral testing. The overexpression of Myl9 and Mustn1 in excitatory neurons of the hippocampus cause hypersocial behavior in WT male mice in the Reciprocal Interaction test **(B)** (p = 0.04) and the Social Preference in the Three Chambers test **(C)** (p = 0.032), with no change in Social Memory in the Three Chambers test **(D)** (p = 0.296). The increased mini EPSC amplitude observed in cKO mice is also mimicked by the overexpression of Myl9 and Mustn1 **(E, F)** (p = 0.039), with no change in mini EPSC frequency **(G)** (p = 0.429). **(H)** Timeline of the acute disruption of actomyosin network by the injection of the ROCK inhibitor Y-27632 (10 mg/kg in saline). Mice were injected *i.p.* with the Rock inhibitor 1 h prior to the first social interaction. **(I)** Y-27632 injection rescued hypersocial behavior observed in cKO mice in the reciprocal social interaction test, without changing sociability in WT mice. Statistics: unpaired t-test (B-G), two-way ANOVA with Tukey post-hoc (I).

The RhoA-associated protein kinase (ROCK) is a critical upstream regulator of actomyosin contractility, e.g., by inhibiting myosin phosphatase thereby leading to enhanced myosin light chain phosphorylation [30], as well as by promoting actin polymerization via activation of Limk1/p-Cofilin [31]. We therefore used the pharmacological inhibitor of ROCK1/ROCK2, Y-27632, to address a causal involvement of activated actomyosin contractility in phenotypes caused by miR379-410 deficiency. Acute treatment (1 h) of rat hippocampal neurons (DIV19) with Y-27632 (10 µM) was able to revert the increase in spine volume caused by simultaneous inhibition of the three miRNA signature with a pLNA cocktail (pLNA1) (Suppl. Fig. 11A-C). In contrast, it had no effect on spine volume in pLNA-control transfected neurons, suggesting that it selectively counteracts elevated actomyosin contractility in response to miRNA inhibition.

Having shown the relevance of ROCK activity for miR379-410 function *in vitro*, we went on to test its role in miR379-410 dependent social behavior in mice *in vi*vo. Intraperitoneal (i.p.) Y-27632 injection (10 mg/kg in saline) in PW15 male mice directly prior (1 - 2 h) to behavioral testing (Fig. 5H) normalized reciprocal social interaction in cKO mice back to WT levels, while no effect was observed in WT mice (Fig. 5H-I). This effect on social behavior was not due to drug-induced changes in locomotor activity (Suppl. Fig. 12). Taken together, these results suggest that ROCK-mediated activation of actomyosin contractility is necessary for hypersociability in adult mice and that this pathway seems to bi-directionally control social behavior in mice.

### MiR379-410 and the muscle actomyosin gene program are deregulated in iNeurons containing an isogenic WBS deletion

Sociability has a high variability in the human population, with a portion of people outside the normal range [4]. While psychiatric disorders (e.g., schizophrenia) and autism spectrum disorders are typically associated with a deficit in social behavior, the opposite trait of hypersociability is exhibited by individuals with specific neurodevelopmental disorders, e.g., Angelman Syndrome and Williams-Beuren Syndrome (WBS). We therefore wondered whether the identified miR379-410-actomyosin pathway might be dysregulated in disorders characterized by hypersociability, focusing on WBS. Induced pluripotent stem cell (iPSC)-derived neurons (iNeurons) represent a valuable cellular system to study the impact of disease-causing mutations at the molecular level. We generated iNeurons harboring a hemizygous genetic deletion at 7q11.23 (7q11 del/+), which is observed in 97% of WBS patients [32]. RNA sequencing from differentiated iNeurons revealed the upregulation of several genes related to actomyosin pathways in 7q11 del/+ versus isogenic WT controls. In fact, the most significant upregulated genesets were associated with the muscle system (Fig. 6A). Specific actomyosin genes found to be strongly upregulated in 7q11 del/+ iNeurons compared to controls overlap with the genes shown to be upregulated in our cKO mice, including ACTA2, MYL9, MYLK and TPM2 (Figure 6B), many of which represent potential direct miR379-410 targets.

**Figure 6:**
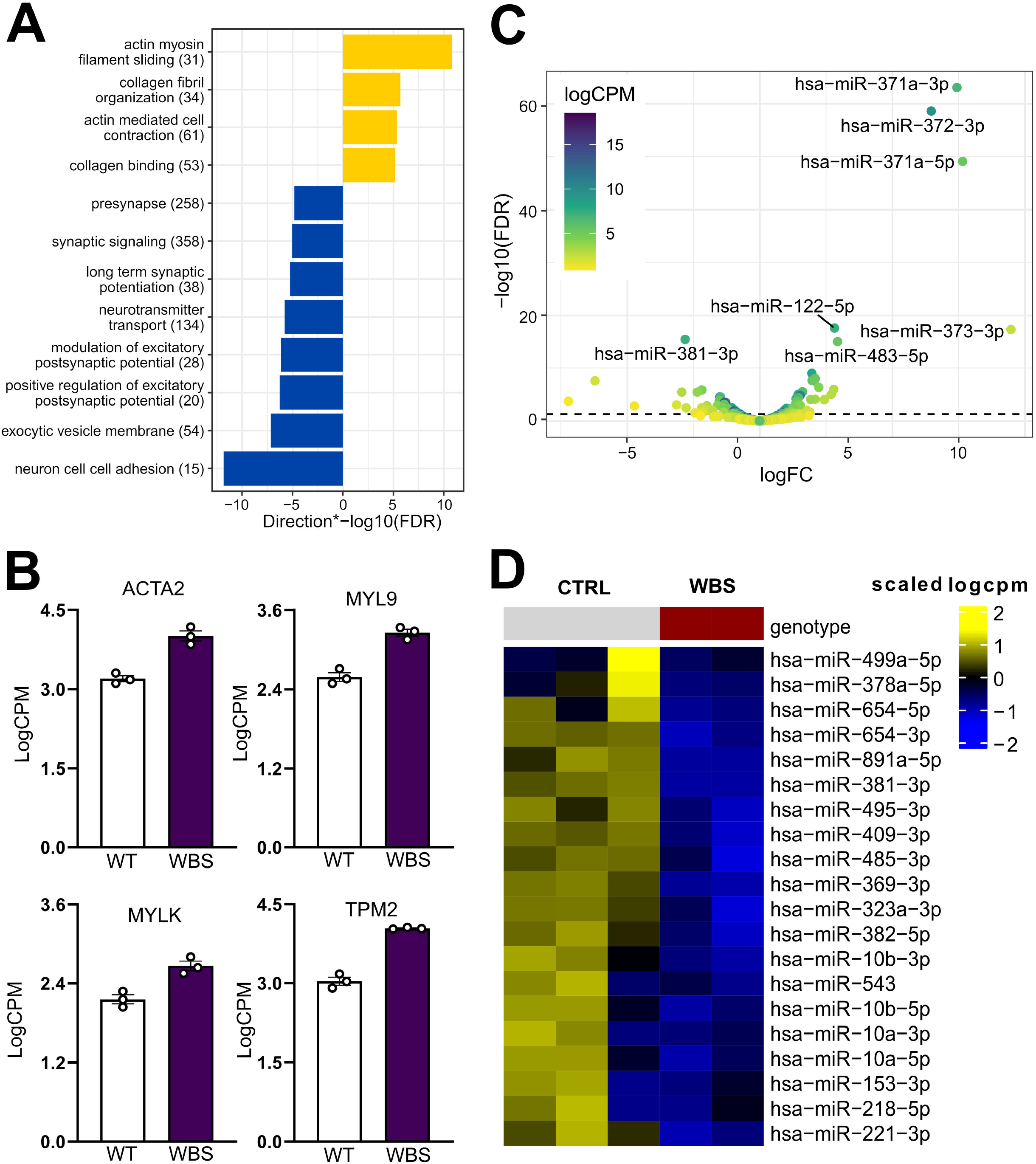
Expression of miR-379-410 miRNAs and actomyosin genes is reciprocally regulated in human iNeurons containing a syntenic WBS deletion. **(A)** GO-term analysis of RNA sequenced from isogenic iNeurons containing a syntenic WBS deletion reveals an upregulation of genes related to actin myosin filament sliding. **(B)** Several mRNAs found to be upregulated in the hippocampus of our cKO mice are also upregulated in iNeurons with a WBS deletion, including Acta2, Myl9, Mylk and Tpm2. **(C-D)** A small-RNA sequencing of WT vs WBS isogenic iNeurons (volcano plot in **(C)** and heatmap showing top 20 downregulated miRNAs in **(D)**) revealed that several members of the miR379-410 cluster are downregulated in WBS iNeurons.

Finally, we asked whether actomyosin gene upregulation in WBS iNeurons might be associated with alterations in miRNA expression. small-RNA sequencing of these isogenic iNeurons samples revealed a differential expression of 99 miRNAs (FDR < 0.05, 56 upregulated, 43 downregulated) (Fig. 6C,D). Strikingly, 13 out of the 43 significantly downregulated miRNAs (FDR<0.05) belong to the miR379-410 cluster, including miR-381-3p, miR-369-3p and miR-495-3p. The downregulation of miR379-410 members concomitant with an upregulation in its actomyosin target gene network in WBS iNeurons is consistent with their functional role as inhibitors of sociability in mice.

## Discussion

In this study, we explored the behavioral effects of a specific deletion of the miR379-410 cluster from excitatory forebrain neurons. By using Emx1-Cre-driven recombination of the miR379-410 allele, we were able to dissociate the hypersociability and anxiety phenotypes that we recently observed in constitutive miR379-410 ko mice [22]. Since Emx1-Cre is almost exclusively expressed in the telencephalic brain regions (olfactory bulb, neocortex, hippocampus), we conclude that the previously observed anxiety [22] likely originated from miR379-410 recombination outside the telencephalon. Future experiments are needed to pinpoint the specific brain regions, e.g. the amygdala, involved in elevated anxiety. At the cellular level, Emx1-Cre is expressed not only in projection neurons, but also in glial cells (astrocytes, oligodendrocytes) [33]. However, we can rule out a contribution of these cell types to behavioral phenotypes, since postnatal deletion of miR379-410 by stereotactic injection of Camk2a-Cre, which is not expressed in glial cell lineages, fully recapitulated hypersociability observed in Emx1-Cre cKO mice. The latter experiment further suggests that the ablation of miR379-410 in adult mice is sufficient to elicit hypersociability, arguing against a major contribution of miR379-410 expressed at embryonic or early postnatal stages. This view is further supported by our observation that social behavior in juvenile Emx1-Cre cKO mice, which underwent recombination alrebehaady at E12.5, was not altered. However, since mature miRNAs turn over slowly, residual miR379-410 expression at early postnatal stages could represent an alternative explanation. Another interesting observation is the striking sex-specific regulation of social behavior by miR379-410. The lack of phenotype in females suggests the existence of female-specific compensatory mechanisms, mediated e.g. by the expression of hormones or X-chromosome encoded gene products.

Our findings from stereotactic brain injections align well with a large body of literature recognizing the importance of the hippocampal formation in the control of social interactions. Our approach, however, does not afford the resolution to draw any conclusions regarding the specific hippocampal subfields involved. As a next step, spatiotemporal expression patterns of the miR379-410/actomyosin network within the hippocampus should be determined using high-resolution miRNA and mRNA in situ hybridization. This data could inform about specific target regions for future circuit manipulations, paving the way for a better understanding of the circuits involved in processing social information.

Gene expression profiling in the hippocampus of miR379-410 deficient mice unraveled a highly specific gene signature with upregulation of a large group of structural actomyosin genes. While many of these genes are typically expressed at high levels in contractile cells, e.g., smooth muscle cells, several of them have been detected by ISH in hippocampal pyramidal neurons *in vivo*, including Myh11, Mylk and Acta2 (Allen brain atlas: mouse.brain-map.org). We further detected expression for several of those genes by qPCR in either FUDR-treated rat primary hippocampal neuron cultures (Myl9; Suppl. Fig. 6C) or in isogenic hIPSC-derived human neurons (Acta2, Myl9, Mylk, Tpm2; Fig. 6). Given the highly selective recombination in principal neurons (i.e., hippocampal pyramidal neurons) using our approach, we consider it highly unlikely that actomyosin gene upregulation is due to miR379-410 inactivation in non-neuronal CNS cells, such as astrocytes or vascular cells (smooth muscle cells, pericytes). Moreover, the miR379-410 precursor RNA Mirg is exclusively expressed by smFISH in pyramidal neurons, but not other cell types present in mixed primary rat neuron cultures (e.g., glial cells) [22], suggesting that miR379-410 deletion, if occurring, should have no effect on the function of non-neuronal cell types. However, even if one postulates pyramidal neuron-specific recombination in the hippocampus, upregulation of actomyosin genes could still occur via a non-cell autonomous mechanism, i.e. involving signaling from neurons to non-neuronal cells. For the following reasons, we consider this alternative mechanism unlikely: first, actomyosin genes are direct targets of miR-379-410 and upregulated in hippocampal neurons of Emx1-Cre cKO mice compared to WT controls (Fig. 3) and second, overexpression of the actomyosin genes Mustn1 and Myl9 in hippocampal neurons using a Camk2a promoter-driven viral construct is sufficient to induce hypersociability (Fig. 5). Together, these results provide strong support for the existence of a miR379-410 regulated actomyosin gene network in mouse hippocampal pyramidal neurons. Our results from pharmacology further demonstrate that the activation of ROCK, an upstream regulator of actomyosin contractility, is required for hypersociability downstream of miR379-410 inhibition (Fig. 5). However, we cannot rule out a contribution of other ROCK targets which are not involved in actin dynamics to the hypersociability phenotype. Moreover, our approach does not allow us to distinguish between actomyosin genes specifically upregulated by miR379-410 deficiency and other non-muscle type myosin present in neurons which have previously been implicated in dendritic spine morphogenesis [34]. More specific approaches, such as the knockdown of specific components of the actomyosin gene network, will be required in this regard.

What could be the physiological significance of actively constraining sociability in adult male mice? It is tempting to speculate that miR379-410-mediated inhibition of actomyosin genes evolved as a mechanism to limit social approach and interaction in polygamous, non-communal species, such as *Mus musculus*. This transition from pro-social to aggressive behavior is observed particularly in males, which have to compete with each other in the search for mating partners [1]. It would be interesting to explore whether miR379-410 and its downstream targets are differentially expressed between communal and non-communal mice, and between males and females.

Finally, we obtained first evidence for the pathophysiological significance of this pathway in the context of WBS. Members of the miR379-410 cluster were strikingly overrepresented among downregulated miRNAs in iNeurons harboring a deletion of the WBS critical region, whereas actomyosin targets were concomitantly upregulated. The latter fits well with recent gene expression network analyses performed in WBS patients [35]. What actually causes the coordinated downregulation of miR379-410 miRNAs in WBS iNeurons is currently unknown. Since loss of the transcription factors Gtf2i/Gtf2ird1 is primarily responsible for hypersociability in WBS patients [36, 37], direct transcriptional regulation of miR379-410 by Gtf2i/Gtf2ird represents a testable hypothesis. In this regard, Gtf2ird1 has been shown to interact with the known miR379-410 upstream regulator Mef2 [38]. Intriguingly, a functional link between the AS gene Ube3a and miR-379-410 has recently been reported in rat hippocampal neurons [21]. This raises the exciting possibility that deregulation of the miR379-410/actomyosin pathway could be a common theme in hypersociability disorders. On the other side of the spectrum, relieving the brake on sociability imposed by miR379-410 could represent a strategy to improve social functioning in disorders encompassing hyposociability, especially since restoring actin dynamics has been reported to be beneficial [39].

## Acknowledgments

We thank Dr. David Colameo for his help with the R scripts used to analyze the dendritic spine data and Cristina Furler for preparing primary rat hippocampal neurons and technical assistance. This project was funded by an SNF project grant (<u>310030_205064</u>; SocioMiR) to G.S and by Ministero della Salute, RC 2019 (ERANET NEURON RRC-2019-2366750 – ALTRUISM) to M.M.

## Author contributions

RN performed the experiments (Fig. 1, 3A-D, 4A-D, 5A-D, and Suppl. Fig. 1, 2, 3, 4, 6A-B, 7, and 10). BL performed the experiments (Fig. 3E-F, 4E-G, 5H-I, Suppl. Fig. 6C, 8, 9, 11, and 12), edited the figures and helped write the manuscript. JW performed electrophysiological recordings (Fig. 2. 5E-G and Suppl. Fig 5). PN sent smallRNAs for sequencing. AMR cloned the luciferase constructs. RF provided technical support. PLG performed bioinformatic analysis of sequencing data and generated related figures (Fig. 3A-D, 6A,C,D, Suppl. Fig. 6A). MM generated WBS isogenic iPSC line and iNeurons (and data were used for Fig. 6). GT provided WBS iPSCs samples for smallRNA sequencing. GS conceived experiments, interpreted results, coordinated collaborative experiments, and wrote the manuscript, which was edited by all authors.

## Data availability

RNA-sequencing data has been deposited to Gene Expression Omnibus (GEO; RNAseq [Fig. 3]: Accession number and Reviewer token available upon request; smallRNAseq [Fig. 6]: Accession number and Reviewer token available upon request). WBS iNeuron data is further available at 7q11.23 Explorer, a web server that allows browsing the data: https://ethz-ins.org/7q/ (password available upon request).

## Materials and methods

### Animals

All animal experiments were performed in accordance with the animal protection law of Switzerland and were approved by the local cantonal authorities (ZH194/21). Mice were housed in groups of 2 - 5 per cage, with food and water ad libitum, and kept in an animal room with inverted light-dark cycle (lights off at 8:30, on at 20:30), and all tests were performed during the dark phase. Mice were handled daily for 5 min for one week before the experiments began. During behavioral experiments, mice were housed individually and allowed to acclimatize to the experimental room in a holding cage for at least 30 min before the beginning of the task. At the end of each experiment, mice were placed back into their original home cage with their littermates. 10 ml/L detergent (Dr. Schnell AG) was used to clean equipment in between trials. Behavioral assays were performed from the least to the most stressful.

### Stereotactic surgery

Anesthesia was quickly induced by 5% isoflurane inhalation in oxygen (1 L/min) and then mice were transferred to a stereotactic frame, having their ears fixed using ear bars, while laying on a warmed plate and wearing a mouth/nose mask with a flow of 2-3% isoflurane in 200 mL/min oxygen. Meloxicam (5 mg/kg in saline) was subcutaneously injected and vitamin A was applied to protect eyes from drying. The mice head was then shaved, cleaned and an incision was made. Bregma and lambda were identified, and the following coordinates were used for the bilateral injections (from bregma): AP -2.06 mm, ML ±1.53 mm and DV -1.51 mm.

The viral vectors (AAV-PHPeB) were produced by the local Viral Vector Facility (VVF9 of the Neuroscience Center Zurich). Mice were injected with 1 µL of virus (per side) using a thin capillary over 5 min, plus 1 additional minute for virus diffusion and then the capillary was removed over 3 min. Local analgesia was provided by suturing the skin while adding Lidocain and Bupivacain (2 mg/kg each) drops on the wound site. Mice were then allowed to recover from the surgery before being group housed again. A second subcutaneous injection of meloxicam (5 mg/kg in saline) was performed 12 h post-surgery, and animals had paracetamol added to their drinking water for the next 48 h. Postoperative health checks were carried out over the 3 days following surgery.

### Behavioral testing

#### Open field

Mice were placed in an open field box (L 45 x W 45 x H 40 cm, TSE System, Bad Homburg, Germany), and allowed to explore it for 10 min. The experiment was videorecorded, and locomotor activity and time spent in the center of the field were automatically analyzed using the TSE VideoMot2 analyzer software (TSE Systems, Bad Homburg, Germany).

#### Reciprocal social interaction test

Mice were single housed for 1 h prior to the start of the test. Two unfamiliar mice from the same genotype (and treatment, in the case of the drug injection cohort) were placed together in the same open field arena (same as the one used for the open field test) and allowed to interact for 10 min. After 1 h kept in single housing, animals underwent the same reciprocal social interaction test with another unfamiliar mouse from the same genotype. The experiment was videorecorded, and the time spent sniffing, scratching and seeking/following each other was quantified by hand.

#### Three chambers test

A three-chambers apparatus was used to test social preference and social memory (L 60 x W 43 x H 22 cm, L 20 cm each chamber). The test consists of three sessions of 10 min each, with 30 min interval between them. On the first session, animals are placed in the center and allowed to explore the arena containing an empty cage (D 8 x H 17 cm) in each side chamber. On the second session (social preference), animals were placed in the center and allowed to explore the arena containing an empty cage in one side chamber and a cage with an unknown mouse (sex- and age-matched) on the other side chamber. On the third session (social memory), animals were placed in the center and allowed to explore the arena containing a cage with the animal used in the previous session in one side chamber and a new unknown mouse (sex- and age-matched) on the other side chamber. The experiment was videorecorded and only the second and third sessions were analyzed by hand, by counting the time spent sniffing each cage. In the social preference, data are presented as time spent sniffing the cage with animal divided by time spent sniffing the empty cage. In social memory, data are presented as time spent sniffing the new unknown mouse divided by time spent sniffing the animal met in the previous session.

#### Elevated plus maze

The elevated plus maze is a test used to measure anxiety-like behavior. Each animal was placed in the center of the elevated plus maze (a cross shaped maze with 2 open arms and 2 closed arms, L 65 x W 5.5 cm each arm, 62 cm elevated from the ground) and their behavior was videorecorded for 5 min. Time spent in the open arms was analyzed by hand. Mice tend to spend more time in the protected closed arms than the open arms. Thus, an increase in the percentage of time spent in the open versus closed arms is interpreted as reduced anxiety behavior.

#### Light-dark box

Mice were placed in a box (L 45 x W 30 x H 25 cm) containing a clear side (L 28.5 cm) and a dark side (L 16.5 cm), with a communicating door and were allowed to explore it for 10 min. The experiment is videorecorded and the number of transitions between the dark and the light box were automatically analyzed using the TSE VideoMot2 analyzer software (TSE Systems, Bad Homburg, Germany).

#### Novel object recognition test

In the first session, mice were placed in an open field arena (L 45 x W 45 x H 40 cm) containing two identical objects placed side by side for 5 min. Mice were then returned to their holding cage for a 15 min break. For the second session, mice were put back in the arena containing the same object from the first session and a novel object for 5 min. The choice of objects was based on a protocol previously published [40] and the objects were randomized between trials and genotypes. The time spent exploring each object in the second session was scored manually and presented as time spent sniffing the novel object divided by time spent sniffing the familiar (from first session) object).

#### Fear conditioning test

Mice were placed in a square open field arena (L 30 x W 30 x H 25 cm) with Plexiglas walls and a metal grid bottom inside the multiconditioning TSE chambers (TSE fear conditioning system, TSE Systems, Germany). In the first session, mice were placed into the conditioning chamber for 3 min, then foot-shocked (2 s, 0.8 mA constant current), and then returned to their home cage. In the second session, after 24 h, mice were placed in the same conditioning chamber for 5 min and percent of time spent freezing was automatically analyzed using the TSE VideoMot2 analyzer software (TSE Systems, Bad Homburg, Germany).

### Primary neuronal cultures

#### Preparation and maintenance of primary neuronal cultures

*In vitro* experiments were approved by the local cantonal authorities (ZH027/21). Primary hippocampal neuronal cultures were prepared from hippocampus or cortex of embryonic day 18 Sprague-Dawley rats (Janvier Laboratories). Cells were plated on acid-treated 13 mm round glass coverslips coated with Poly-L-Lysine (Sigma, P2636) and Laminin (Corning, 734-1098) at a density of 50’000 cells per well on a 24 multi-well plate. Primary cultures were maintained in 500 µL of culture medium (NBP+), containing neurobasal supplemented with B27 (2%), GlutaMAX (1 mM), penicillin (100 U mL^-1^) and streptomycin (100 μg mL^-1^) (all from Invitrogen) in a humidified incubator at 37°C with 5% CO_2_. Cells were fed twice per week by replacing 300 µL of old medium with 325 µL of new warmed culture medium.

#### Primary neuron transfection

Primary cortical (for luciferase experiments) or hippocampal (for dendritic spine experiments) neurons were transfected at in vitro day 7 (DIV 7, cortex) or DIV (hippocampus) with plasmid DNA, miRNA mimics, and/or miRCURY LNA miRNA Power Inhibitors, depending on the experiment. Transfection was performed using antibiotic-free medium (NBP) and Lipofectamine 2000 reagent (Thermo Fisher, 11668019), with a total of 1 µg plasmid DNA (balanced out with pcDNA3 basic vector) for 1 h 30 min. After transfection neurons were washed twice with NBP and treated with 20 μM (2R)-amino-5-phosphonovaleric acid (AP5; R&D Systems, 0105/10), an NMDA receptor antagonist to prevent transfection-induced neuronal overexcitation, diluted in NBP+, for 45 min. Neurons were then washed once in NBP and then returned to the NBP+ (containing half of their NBP+ collected prior to transfection). The plasmids, miRNA mimics and inhibitors used for transfection are listed in Suppl. Table 1 and Suppl. Table 2. For luciferase assays, primary rat cortical neurons were transfected with a WT or mutant 3’UTR-pimirGLO reporter-plasmids, together with 2.5 nM miRNA-134-5p mimic (for Myl) or 2.5 nM miRNA-485-5p mimic (for Tagln) or a scrambled miRNA control. For spine analysis, miRNA overexpression experiment, primary rat hippocampal neurons were transfected with Ctrl-Hp, miR-Hp or miR-Hp2, all encoding for GFP. For spine analysis, pLNA experiment, primary rat hippocampal neurons were transfected with 150 ng GFP and a total of 20 pmol of pLNA control, pLNA 138, pLNA1 cocktail (134-5p, 485-5p and 329-5p, 6.67 pmol each), or pLNA2 cocktail (329-3p, 377-3p and 495-3p, 6.67 pmol each) (Suppl. Table 2).

### iPSC culture, generation of isogenic control and WBS-deletion iPSC lines and iNeuron differentiation

The biological samples from which we derived the iPSC lines used in this study are archived in the Genetic and Genomic Disorders Biobank (GDBank) and the isogenic iPSC lines (control and WBS deletion), have been generated in two consecutive rounds of CRISPR/Cas9 as previously described [32]. In order to obtain cortical glutamatergic neurons (iNeurons), iPSCs were dissociated and plated in Matrigel-coated plates; cells were selected with 1 μg/ml puromycin, to reassure that only the cells with Ngn2-inducible cassette would survive, as previously described in more detail [32].

### RNA extraction

Mouse brain tissue was dissected quickly after cervical dislocation, snap-frozen in liquid nitrogen and kept at -80°C. 1 mL TRIzolTM Reagent (Thermo Fisher, 15596026) was then added, and tissue was homogenized at 4°C in a tissue lyser bead mill (Qiagen) for 2 min at 20 Hz. Subsequently, 200 μL of chloroform (Scharlab, CL02182500) were added and homogenate was incubated for 3 min at room temperature (RT), before being centrifuged at 12’000 × g for 15 min at 4°C. Aqueous phase was transferred to a fresh tube and organic phase was saved for later protein isolation. A second clearing was performed by adding the same amount of chloroform to the aqueous phase and centrifugating. The cleared aqueous phase was then mixed with 2 μL of Glycogen to allow RNA visualization (Ambion, AM9510) and 500 μL of isopropyl alcohol (Sigma, 59304), incubated at RT for 10 min and then centrifugated. RNA pellet was washed twice with 1 mL of 75% ethanol. Lysates were centrifuged for 5 min at 7500 × g at 4°C in between washes. After 30 min drying at RT, pellets were dissolved in 20 μL of nuclease-free H_2_O (Ambion, AM9937) by pipetting up and down. To ensure complete resuspension of the pellet, samples were incubated at 56°C in a heat block. The protocol used for RNA extraction from primary neurons (at DIV 9, 16 and 23) was the same, but only 500 μL of Trizol was used for every 2 wells on a 6-wells plate, and only one chloroform separation step was performed, instead of two. RNA purity and quantity were determined with a Nanodrop 1000 spectrophotometer (DeNovix, Life Science Technologies).

### Poly-A RNAseq analysis

Quantification was performed directly from the reads using salmon 1.3.0 on the GENCODE M20 transcriptome. Genes were filtered for a minimum of 20 reads using edgeR’s (3.32.1) filterByExpr function, and differential expression analysis was performed with edgeR’s quasi-likelihood model [41] including two surrogate variables estimated by SVA [42]. Gene ontology enrichment analyses were performed using the camera pre-ranked method [43] on GO terms with at least 5 quantified genes.

### Short RNA analysis

Short RNA processing was done using sports 1.0 [44] using the authors’ provided annotations. Only reads assigned to miRNAs were kept and summarized at the mature miRNA level. miRNAs were filtered for a minimum of 20 reads using edgeR’s (3.32.1) filterByExpr function, and differential expression analysis was performed with edgeR, including two surrogate variables estimated by SVA [42].

### EnrichMiR and Geneset Enrichment Analysis (GSEA)

We employed our recently developed enrichMiR 0.99.32 [26] to analyze miRNA target enrichment in the set of genes found to be upregulated in the polyA RNA sequencing. Specifically, we used the targetScan 8 target collection (all sites) and the siteoverlap test, with default settings unless stated otherwise. We tested for the miR379-410 members with expression equal or higher than 5 logcpm in the rat brain.

Geneset enrichment analysis (GSEA) was performed using the camera pre-ranked method [43] on the mouse molecular signature database (msigdbr package) collection C5, excluding HPO terms and terms with less than 5 annotated genes in the dataset.

### RT-PCR

Isolated RNA was treated with the TURBO DNase enzyme (Thermo Fisher, AM2238) according to manufacturer’s directions. For detection of mRNAs, DNAse-treated RNA samples were reverse-transcribed using the iScript cDNA synthesis kit (Bio-Rad, 1708891), according to the manufacturer’s recommendations, on a C1000 TouchTM Thermal Cycler (BioRad). Quantitative real-time PCR was performed on a CFX384 Real-Time System (BioRad), using iTaq Universal SYBR Green Supermix (Bio-Rad, 1725121) according to manufacturer’s instructionsFor the detection of miRNAs, DNAse-treated RNA samples were reverse-transcribed using the Taqman MicroRNA Reverse Transcription Kit (Thermo Fisher Scientific). Quantitative real-time PCR was performed using Taqman Universal PCR Master Mix (Thermo Fisher Scientific), according to manufacturer’s instructions. Primers were obtained from Microsynth (mRNA) or Thermo Fisher (miRNA) and their sequences are listed in Suppl. Table 3. Each sample was measured in triplicate. Using the mean Ct-value of these triplicates, real-time RT-qPCR data were analyzed by normalizing each gene of interest to the housekeeping gene U6 (for miRNA), Gapdh or the average of both.

### Cloning

In brief, the mature miRNA-duplexes or a control scrambled sequence with the flanking elements of the miR-30E hairpin required for Drosha processing, were cloned in the 3’UTR of eGFP on a pAAV9-vector under the control of a truncated version of the Camk2a promotor.

Part of the Myl9 (550bp) and Tagln (300bp) 3’UTR (Ensemble transcript IDs: ENSMUST00000088552.7 and ENSMUST00000034590.4) were amplified by touchdown PCR from a mouse brain cDNA library by Phusion Hot Start II DNA Polymerase (Thermo Fisher, F549L) and cloned into the NheI/SalI restriction sites (RS) of the pmirGLO dual-luciferase expression vector (Promega, E1330). Predicted binding-sites of miR-134-5p in Myl9 and miR-485-5p in Tagln were mutated by introducing an SpeI restriction site at the respective miRNA seed match. Myl9 mutation was generated by forward (Fw) and reverse (Rv) primers partially overlapping and amplifying the entire plasmid by PCR from the Myl9-WT. Upon transfection, circularization of the resulting linear product at the complementary overlaps is promoted by inherent bacterial mechanisms. For Tagln, the same primers as for the 3’UTR-amplification and two adjacent to the predicted binding-site, which introduced the SpeI restriction site, were used. The amplified two fragments were digested and ligated at the SpeI site and the resulting insert into the NheI/SalI restriction sites (RS) of the pmirGLO dual-luciferase expression vector. Sequence-verified samples were grown in LB medium overnight at 37°C followed by purification using a MidiPrep Kit (Macherey Nagel) according to the manufacturer’s protocol. Primer sequences are listed in Suppl. Table 3.

### Luciferase assays

After 5 days of expression, cells were lysed and luciferase assay was performed using the Dual-Luciferase Reporter Assay System (Promega, E1910), following a modified protocol previously described [45]. Neurons were washed twice with warm PBS and lysed using 100 μL passive lysis buffer (Promega, E194A) per well (24-well format) for 20 min at 200 rpm. Luciferase activity was measured for 30 μL of cell lysate, transferred into a new 96-well plate, on the GloMax Discover GM3000 (Promega), according to manufacturer’s instructions. The relative luciferase activity was determined by calculating the ratio of Firefly to Renilla signal.

### Immunofluorescence

Primary hippocampal neurons treated or not with FudR and then fixed at DIV 9, 16 or 23 (DIV 7, 14 or 21 after the addition of FudR). Neurons were washed once in prewarmed PBS (Sigma, D8537) and then fixed using 4% PFA/4% sucrose in PBS (Merk, P6148; Sigma, 84097) for 15 min at room temperature. Wells were washed again three times in PBS and then blocked in blocking buffer (10% goat serum and 0.25% Triton in PBS) for 1 h at room temperature. Cells were then incubated with primary antibodies (chicken anti-Map2 [Thermo Scientific, PA1-16751, 1:1000] and rabbit anti-Gfap [Dako, Z0334, 1:500], diluted in blocking buffer) for 1 h at room temperature. After 3 washes with 0.1% Triton/PBS (PBS-T), cells were incubated with secondary antibodies (AlexaFluor 488 goat anti-chicken [Invitrogen, A11039] and Alexa Fluor Plus 647 goat anti-rabbit [Invitrogen, A32733], 1:2000 both, diluted in blocking buffer) for 1 h at room temperature. After 3 washes with PBS-T, cells were incubated with Hoechst 33342 (Thermo Scientific, H3570, 1:2000, in PBS) for 10 min at room temperature. Cells were then washed 3 times with PBS, once with dH_2_O, and coverslips were mounted onto microscope slides using Aqua-Poly/Mount (Chemie Brunschwig, POL18606-20).

### Image acquisition and analysis

Images were acquired with a confocal laser-scanning microscope (CLSM 880, Zeiss) using z-stack (set interval 0.45 μm) with a 63x oil objective at a resolution of 1572 x 1572 pixels, corresponding to an image size of 191.77 × 191.77 μm. Maximum intensity projections of the z-stack images were used for subsequent spine analysis. A total of 10 transfected neurons were imaged experiment and per condition (over two coverslips). Sixteen-bit grayscale images of eGFP (green channel) were analyzed with a custom R-script in the context of the ImageJ-framework. It is freely available via the ImageJ-update site (for more information visit https://github.com/dcolam/Cluster-Analysis-Plugin). In brief, cells are segmented using GFP as a mask. By subsequent skeletonizing and iterating over small branches, tops of spines are identified. Based on these, intensity of the GFP-channel in the selected spine-area and number of spines are measured. Spine volume was calculated by dividing the GFP-intensity in an identified spine by the mean-GFP intensity of the cell (cell body excluded). Spine density was calculated by dividing the number of identified spines by the length of the measured dendrites.

### Electrophysiology

Hippocampal slices from male animals derived from the genetic deletion (WT vs cKO), AAV deletion and Myl9/Mustn1 overexpression cohorts (300 μm thick) were prepared at 4 °C, as previously described [46] (at PW16-20) and incubated at 34°C in sucrose-containing artificial cerebrospinal fluid (sucrose-ACSF, in mM: 85 NaCl, 75 sucrose, 2.5 KCl, 25 glucose, 1.25 NaH_2_PO_4_, 4 MgCl_2_, 0.5 CaCl_2_, and 24 NaHCO_3_) for 30 min, and then held at room temperature until recording.

Whole cell patch clamp recordings were performed at 32 °C on an upright microscope (Olympus BX51WI) under visual control using infrared differential interference contrast optics. Data were collected with an Axon MultiClamp 700B amplifier and a Digidata 1550B digitizer and analyzed with pClamp 11 software (all from Molecular Devices). Signals were filtered at 2 kHz for miniature EPSCs and digitized at 5 kHz. Stimulus evoked postsynaptic currents were filtered at 4 kHz and digitized at 100 kHz. Recording pipettes were pulled from borosilicate capillary glass (Harvard Apparatus; GC150F-10) with a DMZ-Universal-Electrode-Puller (Zeitz) and had resistances between 2 and 3 MΩ.

The extracellular solution (ACSF) was composed of (in mM) 126 NaCl, 2.5 KCl, 26 NaHCO_3_, 1.25 NaH_2_PO_4_, 2 CaCl_2_, 2 MgCl_2_ and 10 glucose. 1 μM TTX and 1 μM Gabazine were added for miniature EPSC (mEPSCs) and cells were recorded at a holding potential of – 70 mV. Synaptic currents for paired-pulse facilitation (inter-stimulus interval: 50 ms) were evoked by mono-polar stimulation with a patch pipette filled with ACSF and positioned in the middle of CA1 stratum radiatum. 1 μM Gabazine was added to isolate AMPA-mediated postsynaptic currents. The intracellular solution was composed of 135 Cs-Gluconate, 5 KCl, 2 NaCl, 0.2 EGTA, 10 HEPES, 4 Mg-ATP, 0.3 GTP and 10 Na_2_-phosphocreatine (adjusted to pH 7.3 with CsOH). Series resistance of CA1 pyramidal neurons (not compensated; range, from 6.5 to 20.3 MΩ; median, 15.3 MΩ; IQR, 4.8 MΩ) was monitored and recordings were discarded if series resistance changed by more than 10%. Membrane potentials were not corrected for liquid junction potential.

### Statistical analysis

All data were analyzed and plotted using GraphPad Prism 9 for Windows (GraphPad Software, San Diego, California USA, www.graphpad.com). The correlation of electrophysiological measures to sociability scores was tested by Pearson correlation. For two-groups comparison, data were analyzed using unpaired two-tailed Student’s t-test. For three-groups comparison, data were analyzed using one-way ANOVA followed by Tukey post-hoc test, when appropriate. For the ROCK inhibition cohort, two-way ANOVA followed by Tukey post-hoc (when appropriate) test was used. Spine data were analyzed using R, linear mixed model fit by REML was applied, t-tests used Satterthwaite’s method [’lmerModLmerTest’], kenward-roger was used as degrees-of-freedom method and dunnettx method for multiple tests was used for p value adjustment. P < 0.05 was considered statistically significant (in graphs, ns p > 0.05, * p < 0.05, ** p < 0.01 and *** p < 0.001). Error bars represent mean ± SEM.

