## Supplemental Figures for "miRNA-mediated inhibition of an actomyosin network in hippocampal pyramidal neurons restricts sociability in adult male mice"

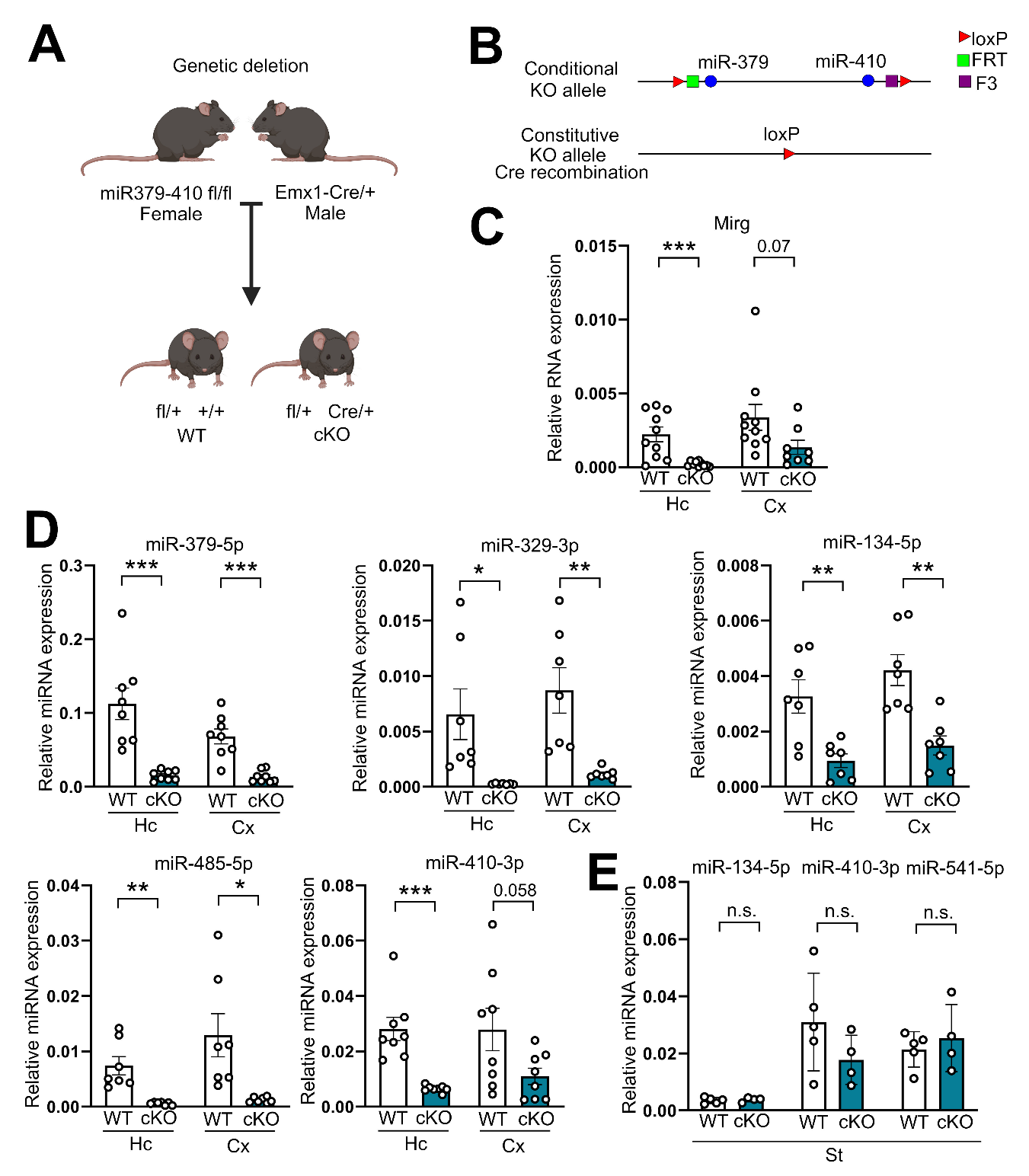

**Suppl. Figure 1:** **Generation and characterization of a mouse line with a conditional deletion of the miR379-410 cluster in excitatory neurons of the forebrain (miR379-410 cko mice).** **(A)** Schematic of the strategy used for the breeding of the animals, in which homozygous miR379-410 flox/flox females were crossed with heterozygous Cre/+ males. Pups born were genotyped as either fl/+ +/+ (WT) or fl/+ Cre/+ (cKO). **(B)** Schematic displaying the sites of flox and the recombination which occur in through Cre recombination. **(C)** The expression levels of Mirg, miR379-410 host gene, are reduced in the hippocampus (Hc, p = 0.0008) and cortex (Cx, p = 0.078) of cKO mice. **(D)** The expression of the mature cluster miRNAs, miR-379-5p (HC p = 0.0005, Ctx p = 0.0001), miR-329-3p (HC p = 0.0176, Ctx p = 0.0033), miR-134-5p (HC p = 0.0038, Ctx p = 0.0013), miR-485-5p (HC p = 0.0014, Ctx p = 0.011) and miR-410-3p (HC p = 0.0002, Ctx p = 0.0589), is reduced in the Hc and Cx of cKO mice. **(E)** The expression of the mature cluster miRNAs, miR-134-5p (p = 0.658), miR-410-5p (p = 0.205) and miR-541-5p (p = 0.535), is not reduced in the striatum (St) of cKO mice. Statistics: unpaired t-test.

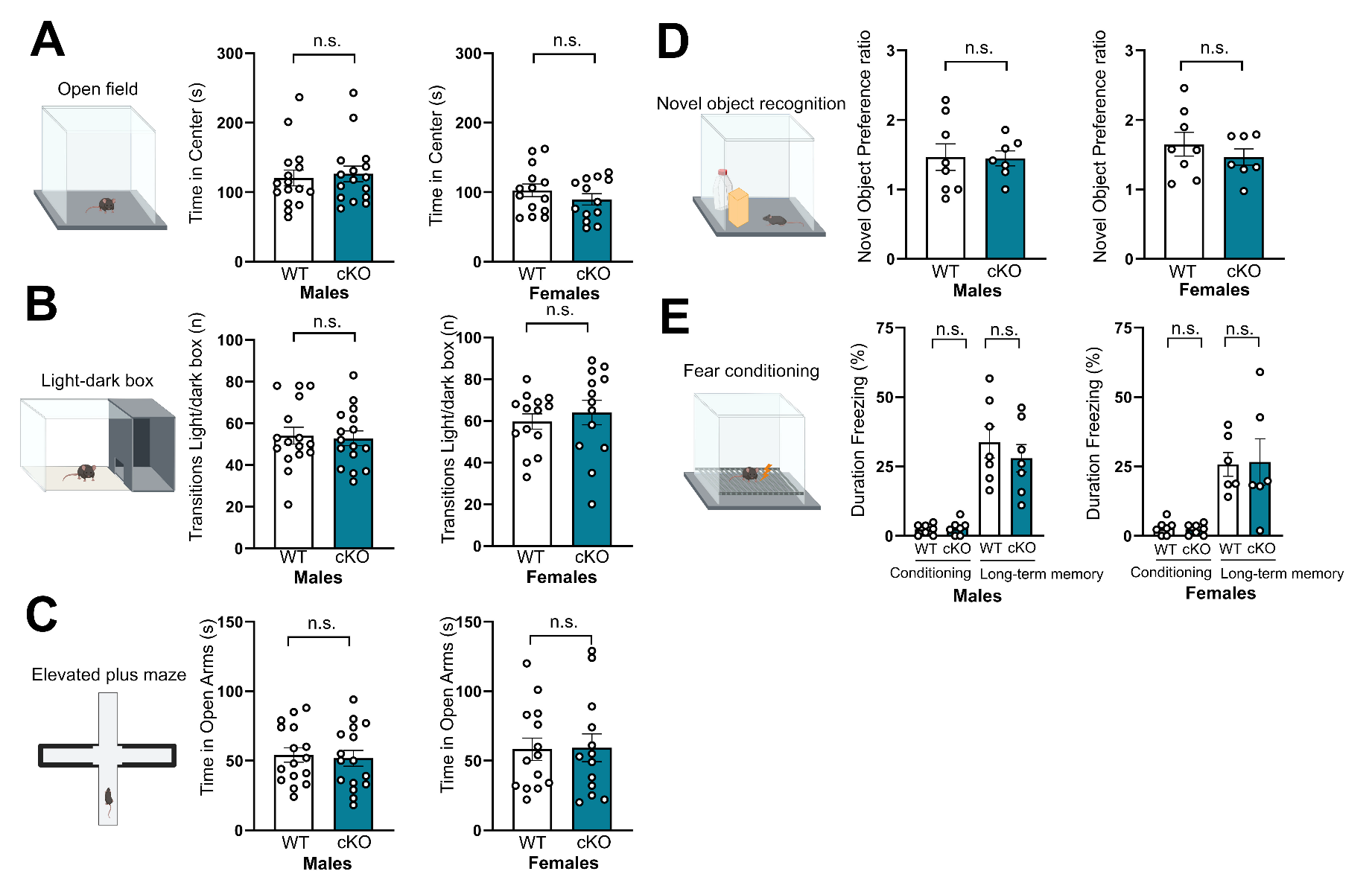

**Suppl. Figure 2:** **Additional behavioral characterization of adult miR379-410 mice.** cKO mice did not display changes in anxiety-like behavior in the open field test **(A)** (males p = 0.715, females p = 0.3), light-dark box **(B)** (males p = 0.817, females p = 0.537), elevated plus maze **(C)** (males p = 0.76, females p = 0.937), and neither did they display changes in memory performance in the novel object recognition test **(D)** (males p = 0.935, females p = 0.407) and fear conditioning test **(E)** (conditioning: males p = 0.632, females p = 0.632, long-term memory: males p = 0.456, females p = 0.93). Statistics: unpaired t-test.

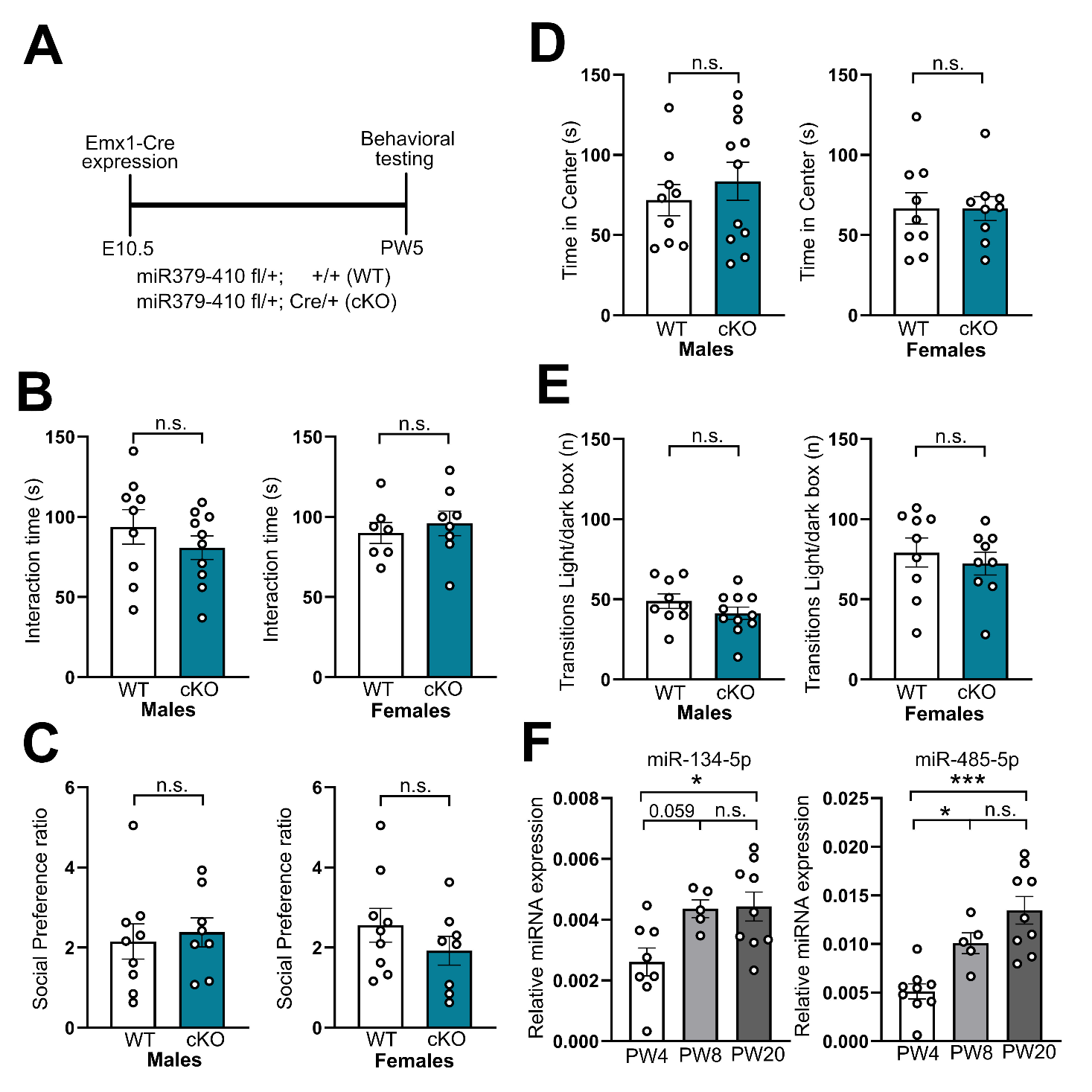

**Suppl. Figure 3:** **Juvenile (PW5) miR379-410 cKO mice did not display any changes in behavior.** **(A)** Timeline of the experiment. Juvenile male and female mice did not display changes in sociability, measured in the reciprocal social interaction test **(B)** (males p = 0.325, females p = 0.569), and in the social preference in the three chambers test **(C)** (males p = 0.7, females p = 0.275). Anxiety-like behavior was also unchanged in the open field test **(D)** (males p = 0.469, females p = 0.99) and light-dark box **(E)** (males p = 0.217, females p = 0.557). **(F)** The levels of expression of miR-134-5p and miR-485-5p (miR-134-5p: PW4 vs PW8 p = 0.0597, PW4 vs PW20 p = 0.019, PW8 vs PW20 p = 0.994 and miR-485-5p: PW4 vs PW8 p = 0.033, PW4 vs PW20 p < 0.0001, PW8 vs PW20 p = 0.176) increase as the animals get older. Statistics: unpaired t-test (B-E) and one-way ANOVA with Tukey post-hoc (F).

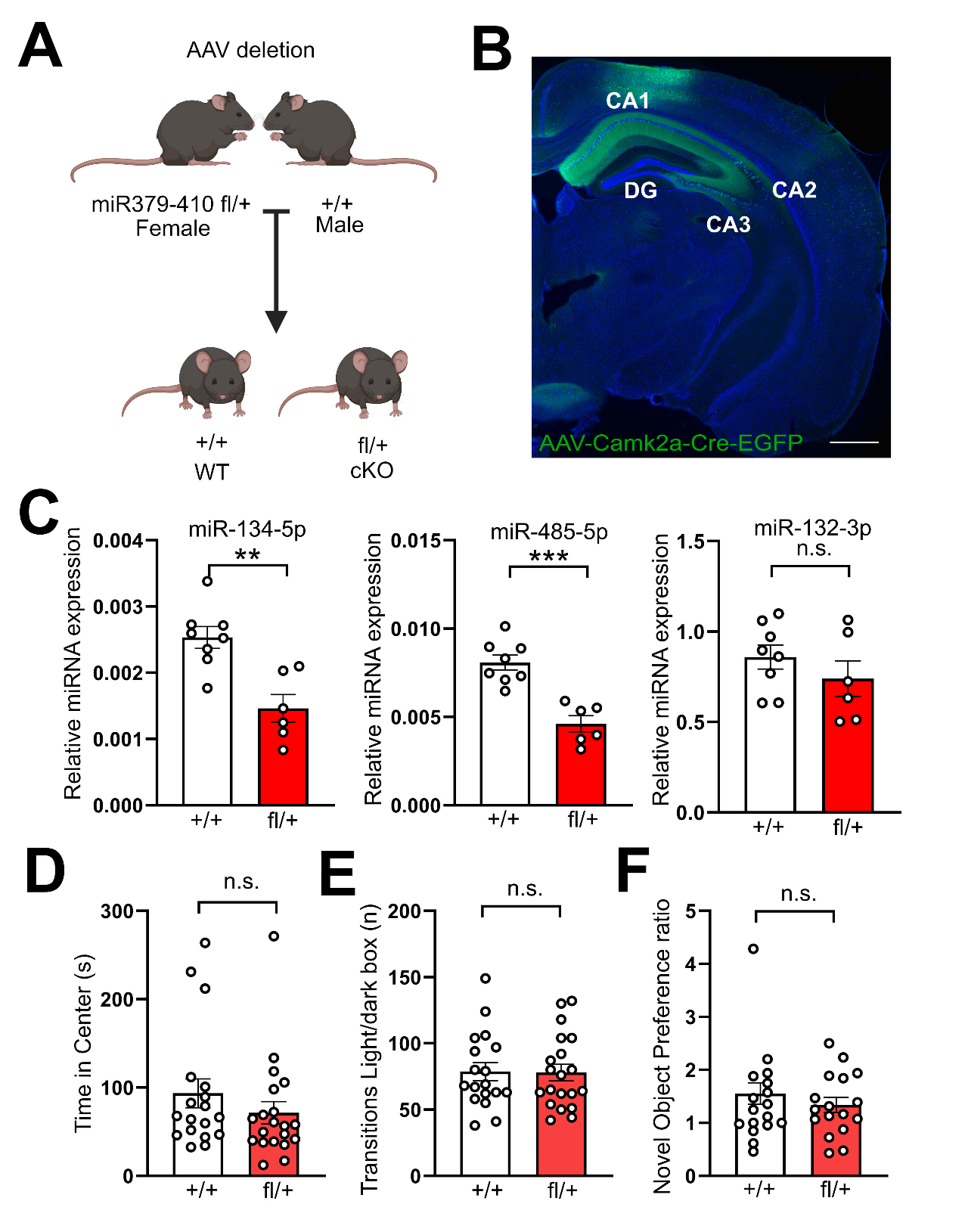

**Suppl. Figure 4: Characterization of male mice with an acute AAV-mediated deletion of the miR379-410 cluster in the hippocampus. (A)** Schematic of the strategy used for the breeding of the animals, in which heterozygous miR379-410 flox/+ females were crossed with WT +/+ males. Pups born were genotyped as either fl/+ (KO after AAV injection) or +/+ (WT). **(B)** Imaging of a brain slice of a mouse injected with the AAV-Camk2a-Cre-GFP demonstrates that the virus is correctly expressed in the CA1, CA2 and CA3 of the hippocampus, with little diffusion to the DG. Scale 500 µm. CA = *cornus ammonius*, DG = dentate gyrus. **(C)** The AAV expression indeed caused a reduction in the expression of two members of the cluster, miR-134-5p (p = 0.0015) and miR-485-5p (p = 0.0001), and did not change the expression of a miRNA unrelated to the cluster, miR-132-3p (p = 0.316). Mice with the adult AAV deletion of the miR379-419 cluster did not display changes in anxiety-like behavior in the open field test **(D)** (p = 0.289) and light-dark box **(E)** (p = 0.938), and neither did they display changes in memory performance in the novel object recognition test **(F)** (p = 0.405). Statistics: unpaired t-test.

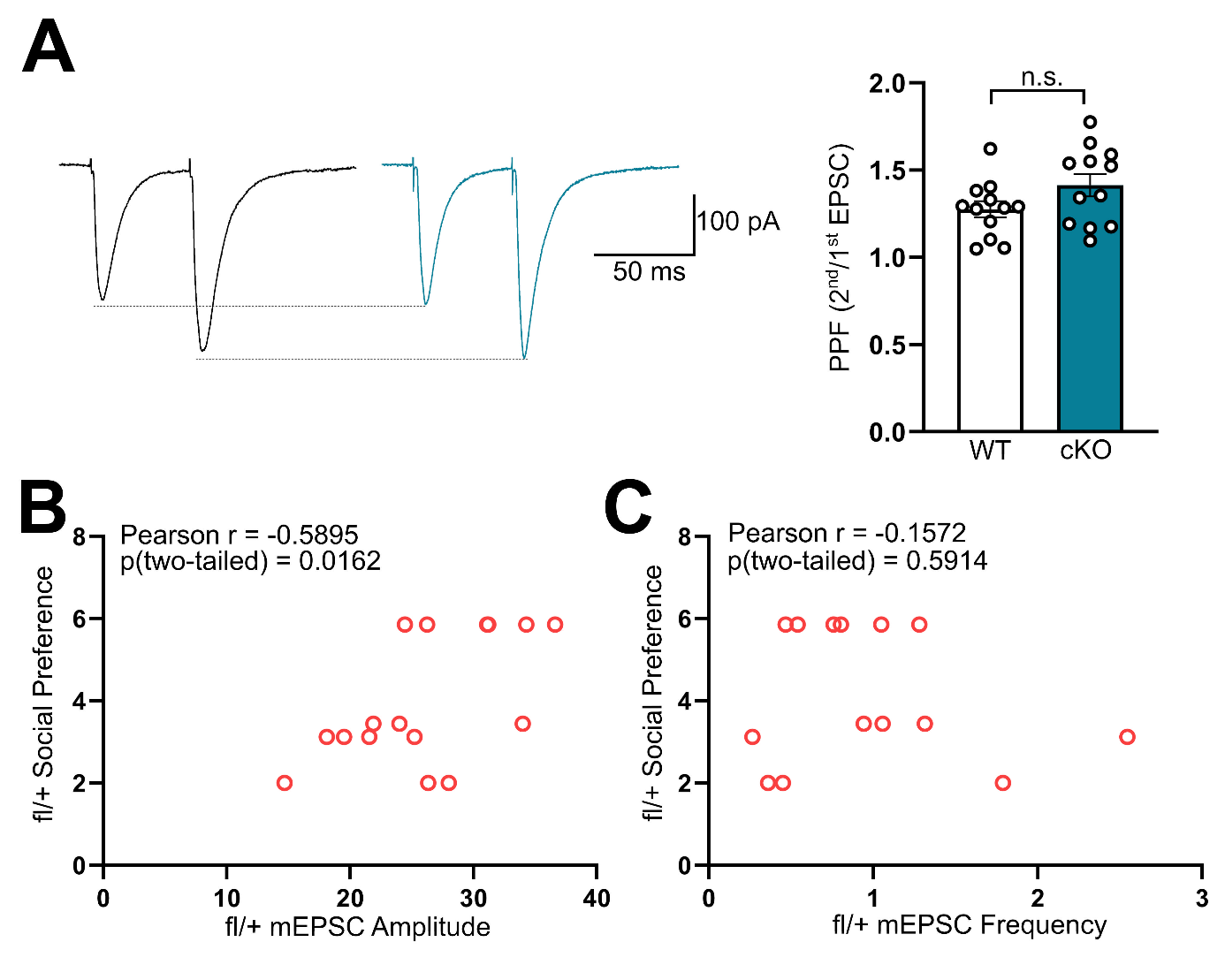

**Suppl. Figure 5 Paired-pulse facilitation was not changed in hippocampal slices of miR379-410 cKO mice. (A)** Paired-pulse facilitation was not changed in cKO mice (p = 0.094). **(B)** mEPSC amplitude in AAV-mediated miR379-410 deletion in hippocampal pyramidal neurons correlates with mice social preference. **(C)** mEPSC frequency in AAV-mediated miR379-410 deletion in hippocampal pyramidal neurons does not correlate with mice social preference. Statistics: unpaired t-test.

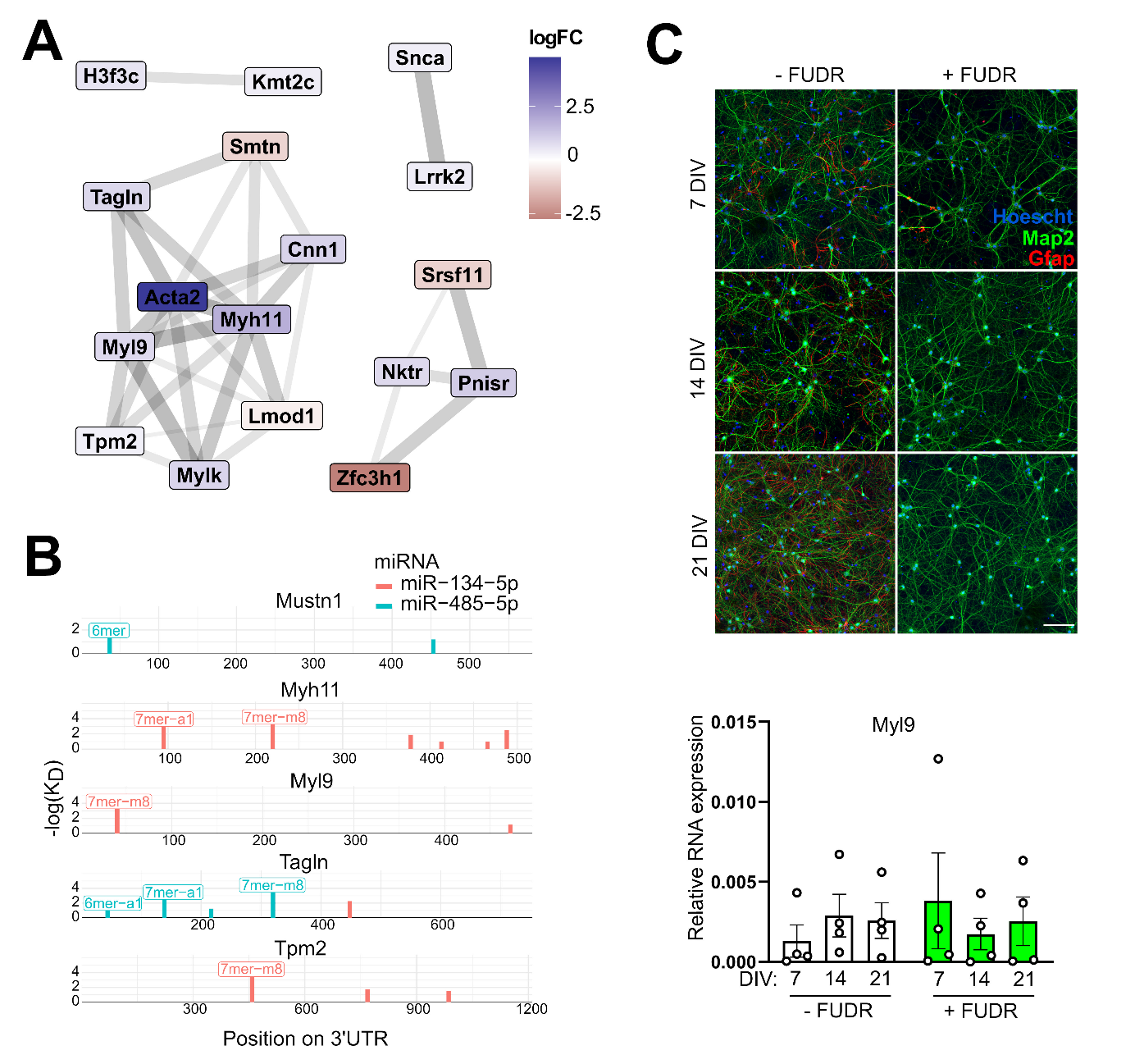

**Suppl. Figure 6:** **Conditional miR379-410 knockout induces an upregulation of actomyosin gene network. (A)** Network of actomyosin genes found to be upregulated in cKO mice. **(B)** ScanMir analysis revealed that the 3’UTR of the actomyosin genes found to be upregulated in cKO mice contain strong binding sites for miR-134-5p and miR-485-5p, members of the three-miRNA signature. **(C)** Primary neurons were treated with FudR (10 µM) from 2 DIV and fixed at 7, 14 and 21 DIV after the addition of FudR and stained for Map2 (as a neuronal marker, green), Gfap (as an astrocytic marker, red) and Hoechst (for nuclei). Scale 100 µm. RT-qPCR of neurons treated or not with FudR revealed that Myl9 is more expressed in neurons over other cells.

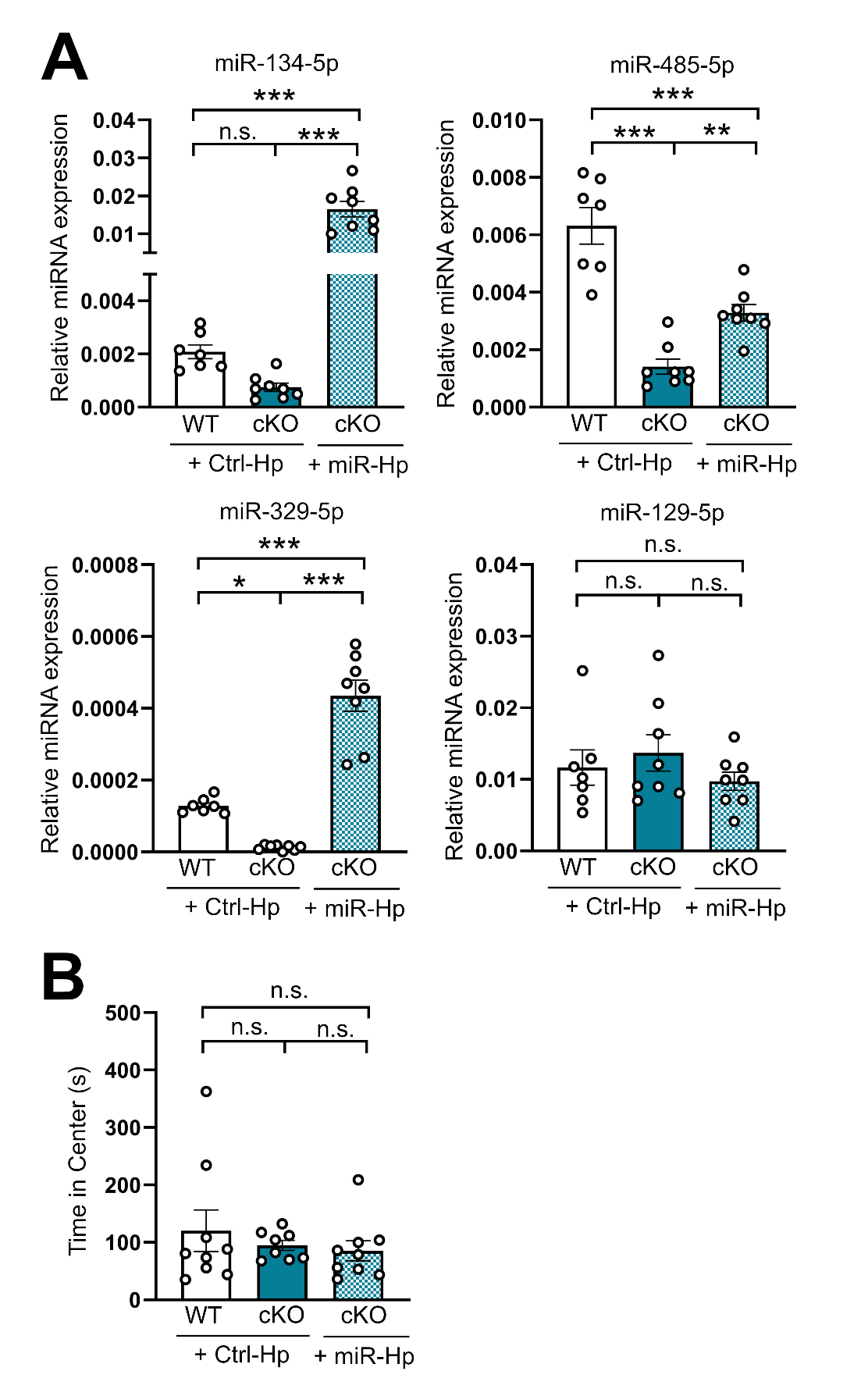

**Suppl. Figure 7:** **Characterization of miR379-410 cKO mice expressing a three-miRNA signature.** **(A)** The expression of the AAV containing hairpins to miR-134-5p (WT Ctrl-Hp vs cKO Ctrl-Hp p = 0.74, cKO Ctrl-Hp vs cKO miR-Hp p < 0.0001, WT Ctrl-Hp vs cKO miR-Hp p < 0.0001), miR-329-5p (WT Ctrl-Hp vs cKO Ctrl-Hp p = 0.017, cKO Ctrl-Hp vs cKO miR-Hp p < 0.0001, WT Ctrl-Hp vs cKO miR-Hp p < 0.0001) and miR-485-5p (WT Ctrl-Hp vs cKO Ctrl-Hp p < 0.0001, cKO Ctrl-Hp vs cKO miR-Hp p = 0.009, WT Ctrl-Hp vs cKO miR-Hp p = 0.0001) induced an increase in the expression of these miRNAs in cKO mice, but no change was observed in the expression of the unrelated miRNA miR-129-5p. **(B)** cKO mice with the re-expression of the three-miRNA signature did not display changes in anxiety-like behavior in the open field test. Statistics: one-way ANOVA with Tukey post-hoc.

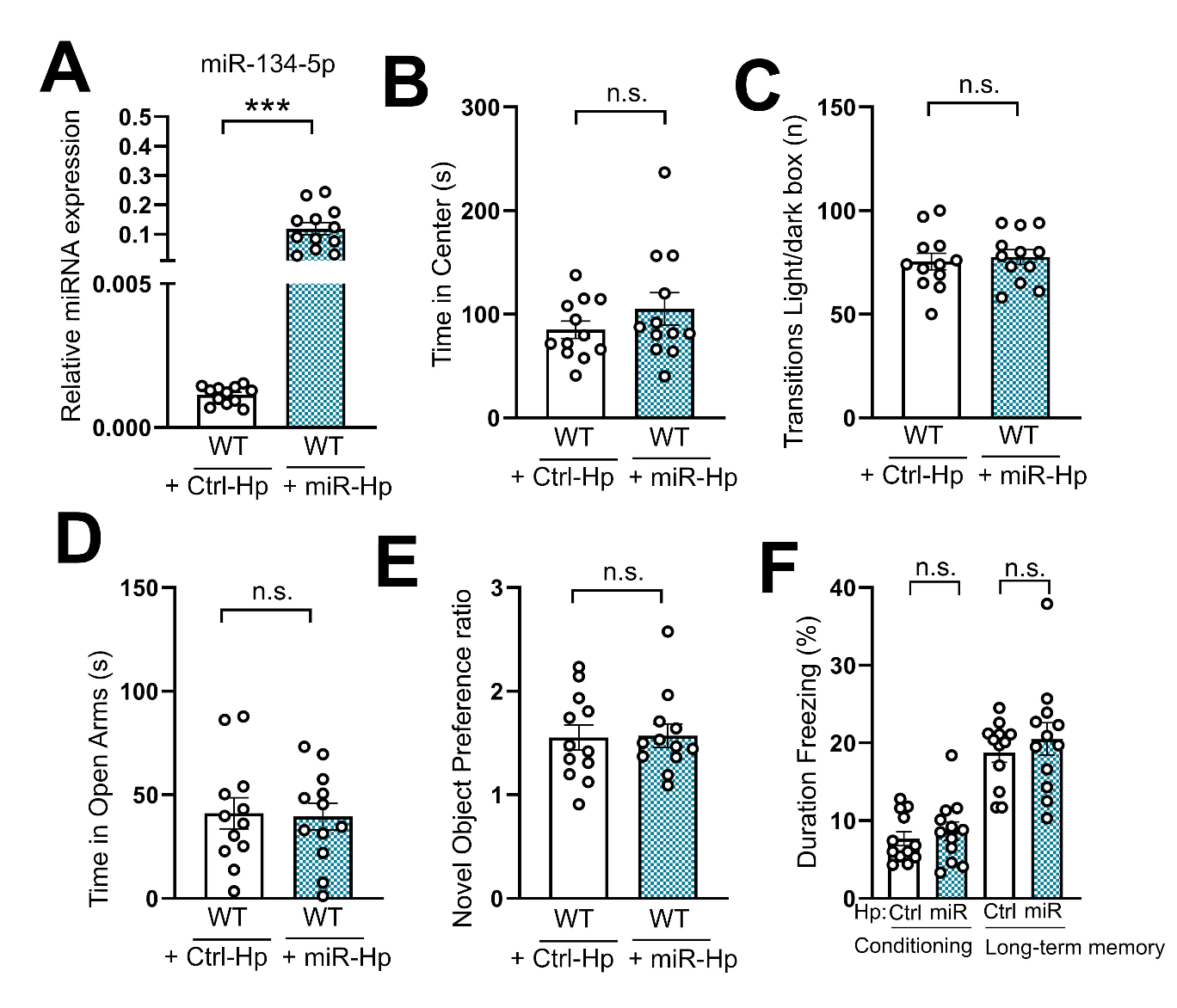

**Suppl. Figure 8:** **Characterization of WT mice overexpressing the three-miRNA signature.** (A) The expression of miR-134-5p was successfully increased (p < 0.0001) after the expression of the miR-Hp AAV. The overexpression of the three-miRNA signature in WT mice did not change anxiety-like behavior in the open field test **(B)** (p = 0.27), light dark box **(C)** (p = 0.68) nor elevated plus maze **(D)** (p = 0.88), and neither did it change memory performance in the novel object recognition test **(E)** (p = 0.92), nor fear conditioning test **(F)** (conditioning p = 0.51, long-term memory p = 0.468). Statistics: unpaired t-test.

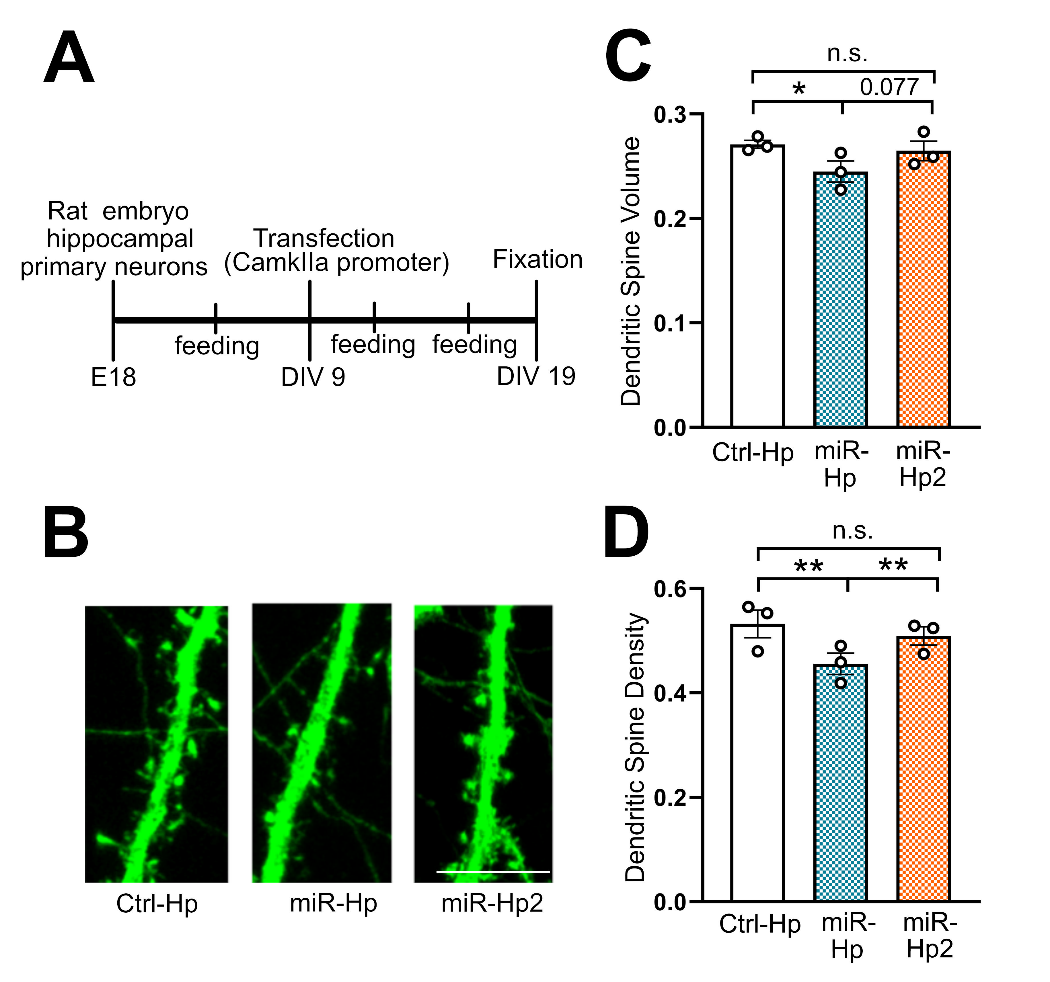

**Suppl. Figure 9: Overexpression of the three miRNA signature reduces dendritic spine volume and density in rat primary neurons. (A)** Timeline of primary neuronal culture and transfection with same miR-Hp plasmid used to produce that AAV from the *in vivo* injections. **(B)** Representative dendritic spines of primary neurons transfected with either Ctrl-Hp, miR-Hp or miR-Hp2 (containing hairpin expressing three other cluster miRNAs, miR 329-3p, 377-3p and 495-5p). Scale: 10 µm. Quantification of spine head revealed that transfection with miR-Hp reduced dendritic spine volume **(C)** and density **(D)** (volume: Ctrl-Hp vs miR-Hp p = 0.034, Ctrl-Hp vs miR-Hp2 p = 0.613, miR-Hp vs miR-Hp2 p = 0.077, volume: Ctrl-Hp vs miR-Hp p = 0.0019, Ctrl-Hp vs miR-Hp2 p = 0.11, miR-Hp vs miR-Hp2 p = 0.007). Statistics: linear mixed model fit by REML, t-tests use Satterthwaite's method ['lmerModLmerTest'], degrees-of-freedom method: kenward-roger, p value adjustment: dunnettx method for 2 tests.

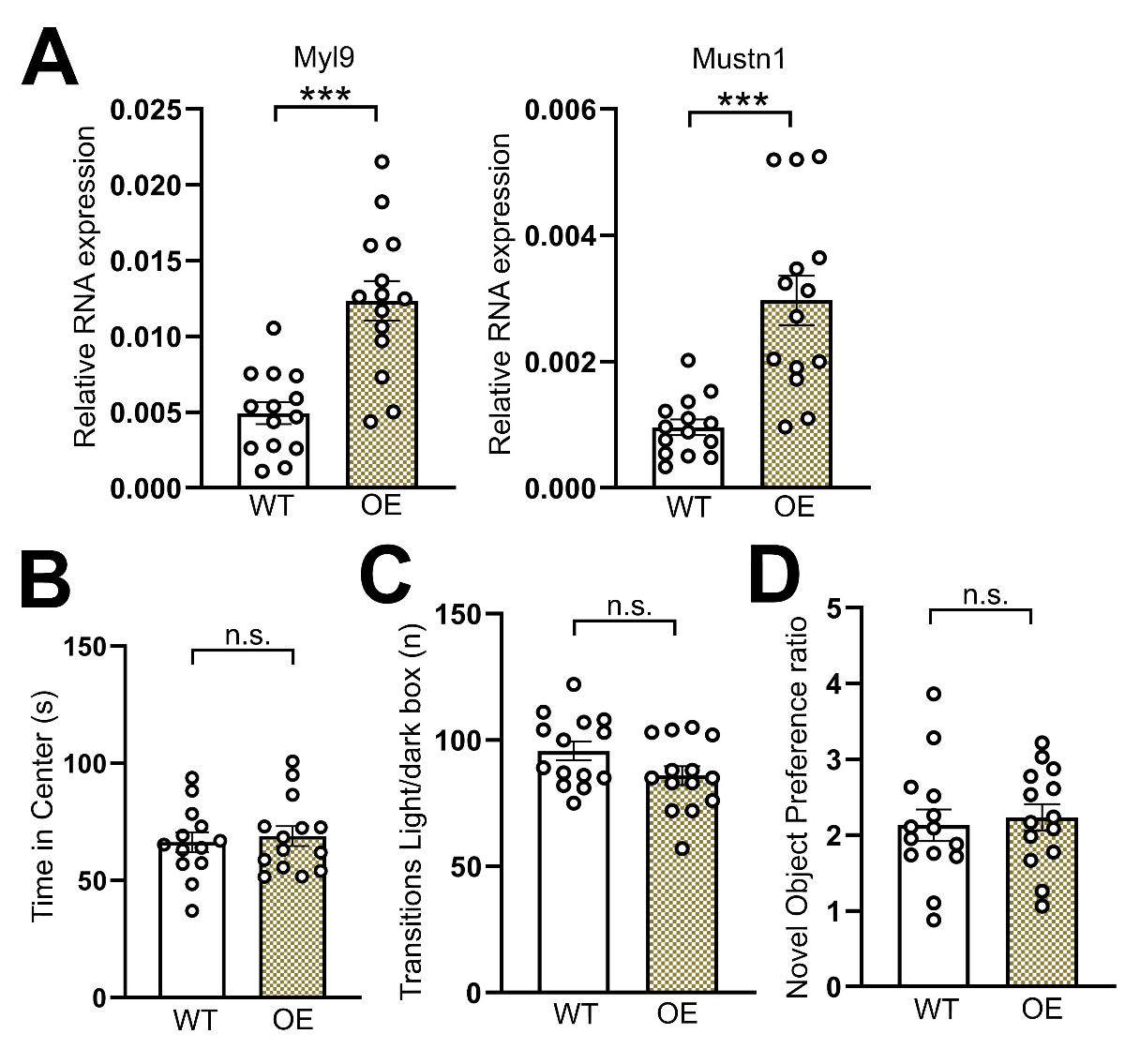

**Suppl. Figure 10:** **Characterization of WT mice overexpressing Myl9 and Mustn1 (OE).** The overexpression of Myl9 and Mustn1 **(A-B)** (p < 0.0001 for both) was confirmed by qPCR dissected from the injected mice. WT mice overexpressing Myl9 and Mustn1 did not display changes in anxiety-like behavior in the open field test **(B)** (p = 0.66) nor light-dark box **(C)** (p = 0.076), neither did they show changes in memory performance in the novel object recognition **(D)** (p = 0.7). Statistics: unpaired t-test.

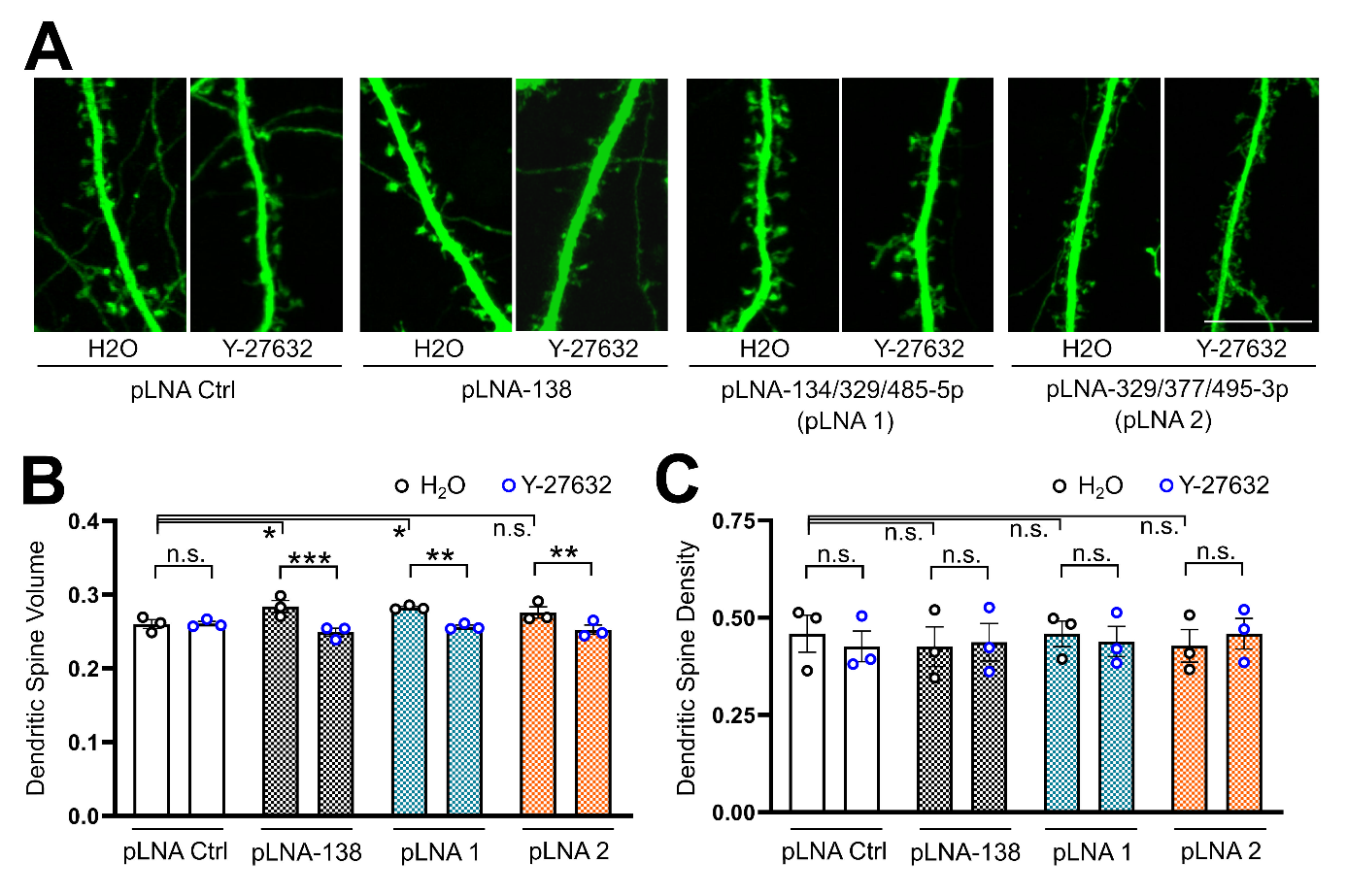

**Suppl. Figure 11: The three miRNA signature increase dendritic spine volume in rat primary hippocampal neurons in a ROCK-dependent manner. (A)** Representative dendritic spines of primary neurons transfected with pLNA Ctrl, pLNA against miR-138 (positive control), pLNAs against the three-miRNA signature (pLNA1, miR-134, -329, and 485-5p), or pLNAs against other three members of the cluster (pLNA2, miR-329, -377, and 495-3p), treated with H_2_O or Y-27632. Scale: 5 µm. **(B)** The transfection of pLNA against miR138 (used as a positive control) and pLNAs against the three-miRNA signature (pLNA1) increased dendritic spine volume, while the transfection of pLNAs against other three members of the cluster (pLNA2, miR-329-3p, miR-377-3p and miR-495-3p) did not change spine volume (pLNA Ctrl vs pLNA-138 p = 0.0168, pLNA Ctrl vs pLNA 1 p = 0.0107, pLNA Ctrl vs pLNA 2 p = 0.115). The treatment with Y-27632 reverts the increased spine volume caused by pLNA-138 and pLNA1 and reduces spine volume in pLNA2 (H_2_O vs Y-27632 in: pLNA Ctrl p = 0.91, pLNA-138 p = 0.0001, pLNA 1 p = 0.0016, pLNA 2 p = 0.0037). **(C)** No pLNA or Y-27632 changes were observed in dendritic spine density. Statistics: linear mixed model fit by REML, t-tests use Satterthwaite's method ['lmerModLmerTest'], degrees-of-freedom method: kenward-roger, p value adjustment: dunnettx method for 4 tests.

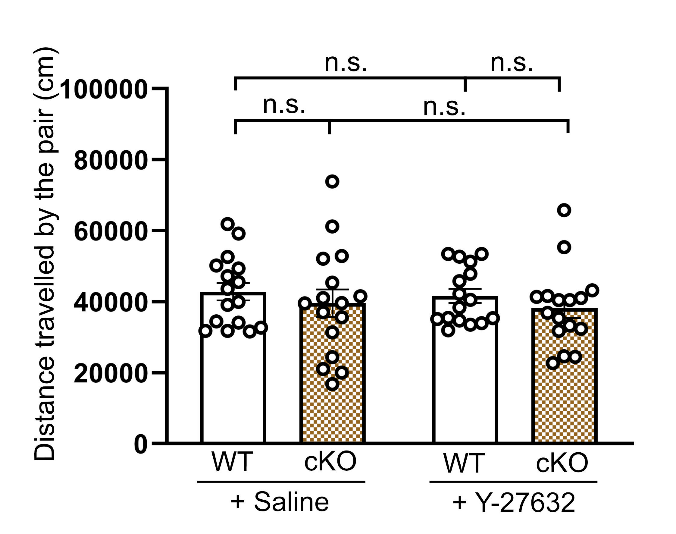

**Suppl. Figure 12: The injection with the ROCK inhibitor did not alter locomotor activity.** Mice intraperitoneally injected with the ROCK inhibitor, Y-27632, did not display changes in locomotor activity, measured by the distance travelled by the pair during the reciprocal social interaction test. Statistics: two-way ANOVA with Tukey post-hoc.

**Supplemental Tables:**

**Suppl. Table 1: Plasmids used for transfection**

|  | Brand | Cat. N° | Concentration |
| --- | --- | --- | --- |
| pcDNA3.1 | Thermo Fisher | V79020 | 850 ng (for pLNA – spines)  900 ng (for miR-Hp – spines)  900 ng (for luciferases) |
| pcDNA3.1-GFP | In-house cloning |  | 150 ng |
| Ctrl-Hp (pAAV[miR30]-Camk2a(short) >EGFP:Scramble[miR30-shRNA]) | VectorBuilder | VB201209-1113kyf | 100 ng |
| miR-Hp (pAAV[3miR30]-Camk2a(short) >EGFP:{mmu-mir-134-5p}:{mmu-mir-329-5p}:{mmu-mir-485-5p}) | VectorBuilder | VB200806-2099dru | 100 ng |
| miR-Hp2 (pAAV[3miR30]-Camk2a(short) >EGFP:{mmu-mir-329-3p}:{mmu-mir-377-3p}:{mmu-mir-495-3p}) | VectorBuilder | VB210106-1160hup | 100 ng |
| pmirGlo Dual Luciferase | Promega | E1330 | 100 ng |

**Suppl. Table 2: Mimics and pLNAs used for transfection**

|  | Brand | Cat. N° | Concentration |
| --- | --- | --- | --- |
| Negative control duplex #1 | AmbionTM | AM17110 | 2.5 nM |
| miR-134-5p duplex | AmbionTM | AM17100-PM10341 | 2.5 nM |
| miR-485-5p duplex | AmbionTM | AM17100-PM10837 | 2.5 nM |
| Negative control pLNA A | Qiagen | 339131-YI00199006 | 20 pmol |
| miR-138 pLNA | Qiagen | 339131-YI04102105 | 20 pmol |
| miR-134-5p pLNA | Qiagen | 339131- YI04104448 | 6.67 pmol |
| miR-329-3p pLNA | Qiagen | 339131- YI04101481 | 6.67 pmol |
| miR-329-5p pLNA | Qiagen | 339131- YI04105741 | 6.67 pmol |
| miR-377-3p pLNA | Qiagen | 339131- YI04104912 | 6.67 pmol |
| miR-485-5p pLNA | Qiagen | 339131- YI04100744 | 6.67 pmol |
| miR-495-3p pLNA | Qiagen | 339131- YI04101229 | 6.67 pmol |

**Suppl. Table 3: Primers RT-qPCR**

| Primer | Sequence |
| --- | --- |
| Mirg | Fw GGCAAGGTCTAGGATGGACA |
|  | Rv CGCCAGCTTCTGAATACTCC |
| Myl9 (rat) | Fw CAGCAGGGAACCCCCAACAC |
|  | Rv AGACATTGGACGTAGCCCTCT |
| Myl9 (mouse) | Fw GGAAGAACCCCACAGACGAG |
|  | Rv CCTCCTCATCAAAGCAGGCA |
| Mustn1 (mouse) | Fw AGGCCCCATCAAGAAGAAGC |
|  | Rv TAGTCCCGCATGACCTGGTA |
| Myl9 (for luciferase) | Fw AAAGCTAGCACTCGCATCCTCAAACACGG |
|  | Rv AAAGTCGACCTGGGGGTGCTGAGACAACT |
| Myl9 (for mutagenesis) | Fw CCATCCACTAGTCCTACCTCCCAACACATACACTCTACCCT |
|  | Rv AGGTAGGACTAGTGGATGGGGTCTAGGCACTGGGGCA |
| Tagln Fw (for luciferase) | Fw AAAGCTAGCCATGACAGGCTATGGGCGAC |
|  | Rv AAAGTCGACCCACTTCTCCCTGCTTACTCC |
| Tagln (for mutagenesis) | Fw AAAACTAGTTAAGGATAGGTGGGAGCTGC |
|  | Rv AAAACTAGTGCCCAGCATCTTACCCCAGAG |
| U6 | Fw AACGCTTCACGAATTTGCGT |
|  | Rv CTCGCTTCGGCAGCACA |
| miR-129-5p | Taqman Catalog No: 4427975 / 000590 (Thermo Fisher) |
| miR-132-3p | Taqman Catalog No: 4427975 / 000457 (Thermo Fisher) |
| miR-134-5p | Taqman Catalog No: 4427975 / 001186 (Thermo Fisher) |
| miR-329-3p | Taqman Catalog No: 4427975 / 000192 (Thermo Fisher) |
| miR-329-5p | Taqman Catalog No: 4427975 / 463347_mat (Thermo Fisher) |
| miR-379-5p | Taqman Catalog No: 4427975 / 001138 (Thermo Fisher) |
| miR-410-3p | Taqman Catalog No: 4427975 / 001274 (Thermo Fisher) |
| miR-485-5p | Taqman Catalog No: 4427975 / 001036 (Thermo Fisher) |
| miR-541-5p | Taqman Catalog No: 4427975 / 002562 (Thermo Fisher) |
| U6 snRNA | Taqman Catalog No: 4427975 / 001973 (Thermo Fisher) |
